# Spatial Transcriptomics Reveals the Requirement of ADGRG6 in Maintaining Chondrocyte Homeostasis in Mouse Growth Plates

**DOI:** 10.1101/2023.09.21.558739

**Authors:** Fangzhou Bian, Victoria Hansen, Hong Colleen Feng, Yanshi Chen, Ryan S. Gray, Chia-Lung Wu, Zhaoyang Liu

**Affiliations:** Center for Craniofacial Molecular Biology, Herman Ostrow School of Dentistry, University of Southern California, Los Angeles, CA 90033, USA; Center for Musculoskeletal Research, Department of Orthopedics and Rehabilitation, University of Rochester Medical Center, Rochester, NY 14642, USA; Department of Biology, University of Rochester, Rochester, NY 14642, USA; Department of Nutritional Sciences, Dell Pediatric Research Institute, The University of Texas at Austin, Austin, TX 78723, USA; Department of Orthopaedic Surgery, Keck School of Medicine, University of Southern California, Los Angeles, CA 90033, USA

**Author notes:** Author for correspondence Zhaoyang Liu, 2250 Alcazar Street, Los Angeles, CA 90033, USA. co-senior authors (Chia-Lung Wu, Zhaoyang Liu).

**Keywords:** ADGRG6/GPR126, Spatial Transcriptomics, Chondrocytes, Homeostasis, Growth Plate, GPCR, Formalin-fixed Paraffin-embedded (FFPE) Tissues, Postnatal Cartilage

## Abstract

The growth plate is essential for maintaining skeletal growth; however, the mechanisms governing postnatal growth plate homeostasis are poorly understood. Here we show that ADGRG6/GPR126, a cartilage-enriched G protein-coupled receptor (GPCR), is dispensable for embryonic limb development but is required for postnatal growth plate homeostasis. *Adgrg6* ablation in osteochondral progenitor cells or postnatal chondrocytes leads to reduced cellularity and impaired maintenance of the resting zone in the growth plate, coupled with increased cell death and reduced cell proliferation. *Adgrg6* mutant growth plates also exhibit disorganized extracellular matrix structures and dysregulated hypertrophic differentiation. Furthermore, using a novel spatial transcriptomics workflow that applies to FFPE tissue sections of mineralized mouse knee joints, we demonstrate that *Adgrg6* ablation leads to reduced SOX9 expression, induced Indian hedgehog (IHH) signaling, and a precocious chondrogenic-to-osteogenic conversion of the growth plate chondrocytes that may be driven by increased POSTN/integrin receptor signaling. We further demonstrated that ADGRG6 regulates the proper formation of the resting zone growth plate by maintaining the PTHrP and SOX9-positive cell populations. Altogether, our findings elucidate the essential role of ADGRG6 in maintaining chondrocyte fate, survival, and homeostasis of the postnatal growth plates.

## Introduction

The cartilage growth plate plays an essential role in postnatal linear growth of the long bone ^1, 2^. In humans, growth plates emerge around birth and maintain skeletal growth until closing at puberty. Trauma, genetic mutations, and other conditions that cause skeletal abnormalities can lead to growth plate defects and accelerate growth plate closure ^3^. However, the molecular mechanisms that regulate postnatal growth plate homeostasis remain poorly understood.

The growth plate consists of distinct zonal layers of chondrocytes, called resting, proliferative, and prehypertrophic/hypertrophic chondrocytes ^1^. The resting zone chondrocytes provide precursors for the adjacent columnar chondrocytes in the proliferative zone ^4, 5^. They highly express the parathyroid hormone-related protein (PTHrP, also known as PTHLH), which keeps the chondrocytes proliferating and delays the production of Indian hedgehog (IHH). Conversely, IHH released from the prehypertrophic and early hypertrophic chondrocytes promotes the expression of PTHrP. This PTHrP/IHH feedback loop helps to ensure chondrocyte proliferation and prevent premature hypertrophy ^1, 6, 7^. While some hypertrophic chondrocytes undergo apoptosis after terminal differentiation, lineage-tracing studies have provided evidence that many of these cells convert into osteoblasts/osteocytes, marrow stroma cells, and adipocytes ^4, 8, 9, 10^. Recent studies have shown that SOX9 is an important regulator of osteogenic plasticity of growth plate chondrocytes ^11^. Ablation of *Sox9* in postnatal cartilage drives growth plate chondrocytes to a rapid conversion of osteoblast-expressing pattern ^11^. Together, the development and maintenance of the growth plate are mediated by multiple regulators and synchronized signaling pathways, so further research is required to fully encode the mechanisms that control growth plate homeostasis.

Chondrocytes can respond to various external signaling via the transmembrane G protein-coupled receptors (GPCRs), the largest class of membrane proteins in the human genome ^12^. The classical GPCR-mediated signaling transduction is dependent on G proteins, which are classified into four major families according to their α subunit (Gs, Gi/o, Gq/11, and G12/13). Different G protein families can direct diverse signaling outcomes. For example, Gs signals through adenylyl cyclase to generate cyclic adenosine monophosphate (cAMP) and activate Protein Kinase A (PKA) and cAMP Response Element-Binding Protein (CREB), while Gq/11 signals through beta-type phospholipase C (PLC-β) to hydrolyze PIP2 to IP3 and result in induced calcium release and activation of Protein Kinase C ^13, 14, 15^. The most well-studied GPCR in cartilage is PTH1R (also known as PPR), a receptor for parathyroid hormone (PTH) and PTHrP, all of which are well-documented to regulate cartilage and bone development ^16, 17, 18, 19, 20, 21, 22^. PTH1R can signal through both Gs and Gq/11 to regulate the quiescence and proliferation of the stem cells/chondroprogenitors in the resting zone of the growth plate ^22, 23^. Thus, GPCRs and G protein-mediated signaling are required in growth plate homeostasis, and different G-protein stimulatory families may be responsible for distinct biological functions ^24^.

ADGRG6 (Adhesion G Protein-Coupled Receptor G6, previously called GPR126) is highly enriched in cartilaginous tissues, modeling spine and ribcage deformities in *Adgrg6* conditional mutant mice ^25, 26, 27^. We and others have shown that ADGRG6 can signal through Gs via the cAMP/PKA/CREB axis in various cell populations ^25, 28, 29^. However, the biological function and molecular mechanisms of ADGRG6 in growth plate homeostasis are still largely unknown. To address these questions, we show here that *Adgrg6* is dispensable for embryonic limb development but is required for postnatal growth plate homeostasis. Using genetic mouse models and spatial transcriptomics approaches, we found that loss of *Adgrg6* in osteochondral progenitor cells and postnatal chondrocytes leads to precocious chondrogenic-to-osteogenic conversion and increased cell death, associated with ectopic upregulation of catabolic enzymes and dysregulated hypertrophic differentiation. These findings provide new insights into the role of a cartilage-enriched GPCR in maintaining chondrocytes homeostasis of the postnatal growth plates.

## Results

### Loss of *Adgrg6* in osteochondral progenitor cells results in growth plate defects in postnatal mice

To investigate the role of *Adgrg6* in the growth plate, we crossed mice harboring an *Adgrg6^f/f^* conditional allele ^28^ to the *Col2a1Cre* mouse strain, which recombines in osteochondral progenitor cells ^30^. At postnatal (P) day 1, skeletal preparations revealed no apparent defects in the size and patterning of the skeletons in the *Col2a1Cre; Adgrg6^f/f^* mutant mice (called *Adgrg6* cKO hereafter) compared with the Cre (-) littermate (wild-type) controls (Fig. 1A, B). Histological analysis with control and mutant tibia epiphyses showed comparable morphology at P1 (Fig. 1C, D), albeit a mild disorganization of the hypertrophic cells in the *Adgrg6* mutant growth plates (black segment, Fig. 1D’). Immunohistochemistry (IHC) analysis confirmed the reduced expression of ADGRG6 in the mutant epiphyses (Supplementary Fig. 1A-C). These data indicate that *Adgrg6* is dispensable for embryonic limb development.

**Figure 1.**
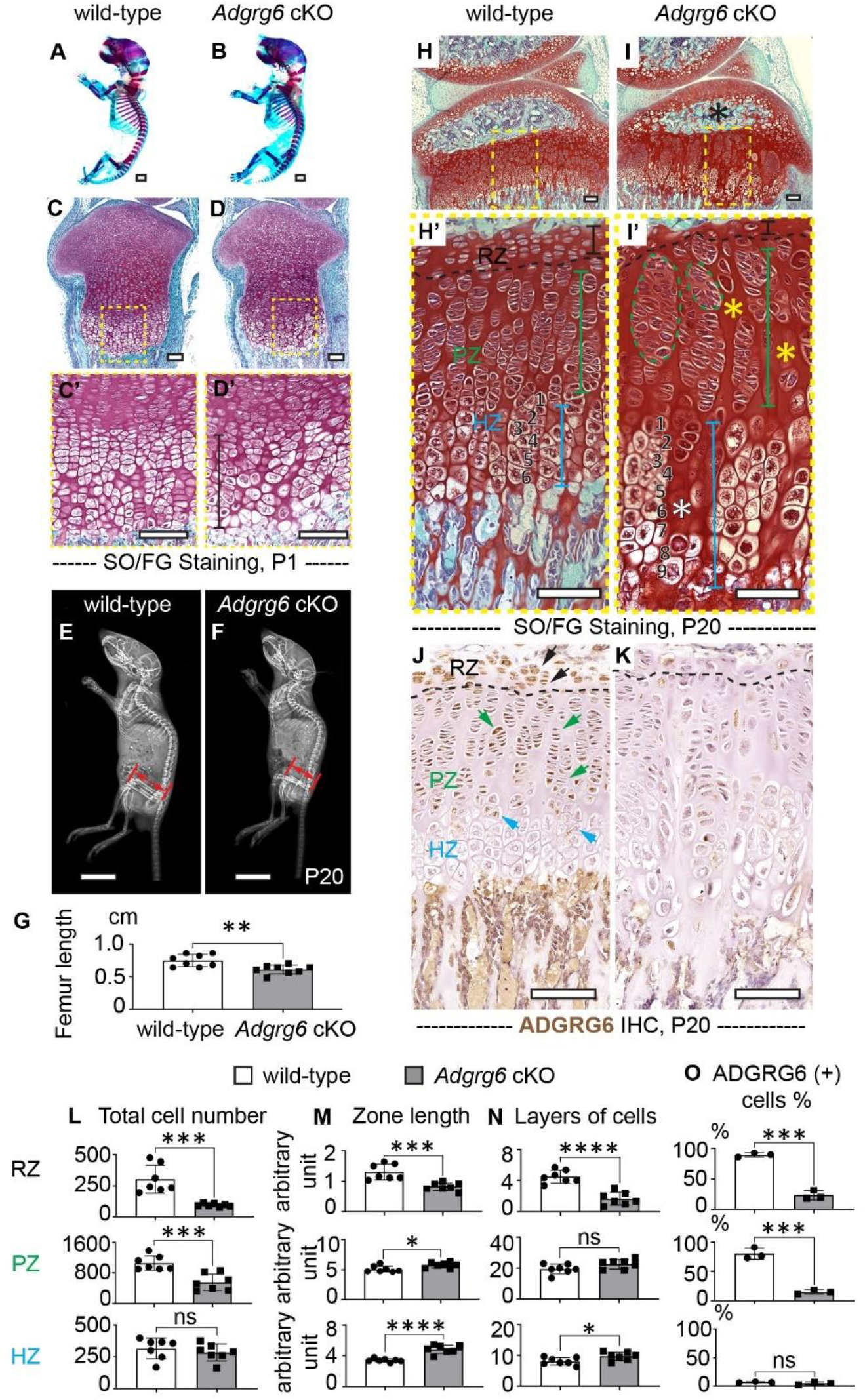
Loss of *Adgrg6* in osteochondral progenitor cells results in growth plate defects in postnatal mice. **(A, B)** Skeletal preparations of the wild-type and *Adgrg6* cKO mice at P1. (n=3 per genotype.) **(C-D’)** SO/FG staining of the control and mutant tibias at P1. The mild disorganization of the hypertrophic cells in the mutant growth plate is labeled with a black segment. (n=3 per genotype.) **(E-G)** X-ray analysis of P20 control and mutant mice. The length of the femur is indicated with double-arrowed segments in E and F and is quantified in G. (n=8 per genotype. ** p=0.0042.) **(H-I’)** SO/FG staining of the control and mutant tibia growth plates at P20. The impaired formation of the secondary ossification center is indicated with a black asterisk. The resting zones are indicated with black dashed lines and segments. The proliferative and hypertrophic zones are indicated with green and blue segments. The formation of acellular clefts in the mutant proliferative and hypertrophic growth plates are indicated with yellow and white asterisks. Cells formed into clusters are circled with green dashed lines. The layer of cells in a hypertrophic cell column is labeled in H’ and I’. (n=7 per genotype.) **(J, K)** IHC analysis of ADGRG6 in control and mutant mice at P20. ADGRG6 (+) cells in the resting, proliferative, and hypertrophic zones are indicated with black, green, and blue arrows, respectively. (n=3 per genotype.) **(L-O)** The total cell number, zone length, and layers of cells in each zone are quantified in L-N. (n=7 per genotype.) The percentage of ADGRG6 (+) cells is quantified in O. (n=3 per genotype.) (L: *** p=0.0003 for RZ; p=0.0006 for PZ; ns, p=0.4652 for HZ. M: *** p=0.0008 for RZ; *p=0.0213 for PZ; **** p<0.0001 for HZ. N: **** p<0.0001 for RZ; ns, p=0.0818 for PZ; *p=0.0303 for HZ.) Bars are plotted by means ± SD, two-tailed t-test. SO/FG: Safranin O/Fast Green. IHC: Immunohistochemistry. RZ: resting zone; PZ: proliferative zone; HZ: hypertrophic zone. Scale bar: 1mm in A and B; 1cm in E and F; 100µm in C-D’ and H-K.

However, at P20, X-ray analysis showed growth retardation of the *Adgrg6* cKO mice, exhibiting a shortening of the appendicular skeleton relative to the controls (Fig. 1E-G). Histological analysis revealed a 100% penetrance of a disorganized growth plate phenotype in the mutant tibias, associated with an impaired formation of the secondary ossification center (Fig. 1H-I’) and reduced cellularity in the resting and proliferative growth plates (Fig. 1H-I’, L). The length of the mutant resting zone is decreased by approximately 50%, with only 1-3 layers of cells retained (Fig. 1I’, M, N). Cells in the mutant proliferative growth plates tend to form clusters instead of columns (green circles, Fig. 1I’). The extracellular matrix (ECM) of the mutant growth plate is disorganized as evidenced by acellular clefts throughout the proliferative and hypertrophic zones (yellow and white asterisks, Fig. 1I’). Meanwhile, *Adgrg6* cKO mice exhibit elongated prehypertrophic/hypertrophic growth plates (Fig. 1I’, M, N). Reduced cellularity and increased chondrocyte hypertrophy were also observed in the articular cartilage of the *Adgrg6* cKO mice at P20 (Supplementary Fig. 1D-E’). When these mice reached adulthood (8 months), they displayed more severe loss of cellularity, reduced proteoglycan staining, and disorganized ECM; however, a complete growth plate closure was not observed (Supplementary Fig. 1F, G).

IHC analyses showed that ADGRG6 is highly expressed in the resting and proliferative chondrocytes of the control growth plates, while its expression is largely depleted in the *Adgrg6* mutants (Fig. 1J, K, O). Interestingly, ADGRG6 exhibits a pattern of expression that is complementary to that of PTH1R, which is highly enriched in the prehypertrophic/hypertrophic chondrocytes and osteoblasts, but with lower expression in the proliferative chondrocytes^1^ (Supplementary Fig. 1H, I). Altogether, these findings suggest that ADGRG6 does not affect embryonic limb development but regulates the homeostasis of the postnatal growth plates.

### Establishment of a spatial transcriptomics workflow on the postnatal knee joint

To map the molecular regulatory mechanisms of *Adgrg6* in postnatal growth plates, we established a novel spatial transcriptomics workflow that applies to formalin-fixed, paraffin-embedded (FFPE) tissue sections of the mineralized mouse knee joints (see Materials and Methods for details). We embedded one control and one *Adgrg6* cKO mutant P20 hind limb into a single paraffin block to reduce the batch-to-batch discrepancy (Fig. 2A, B). After sequence alignment and image mapping, this workflow detected 19,297 genes with 1,852 spatial spots.

**Figure 2.**
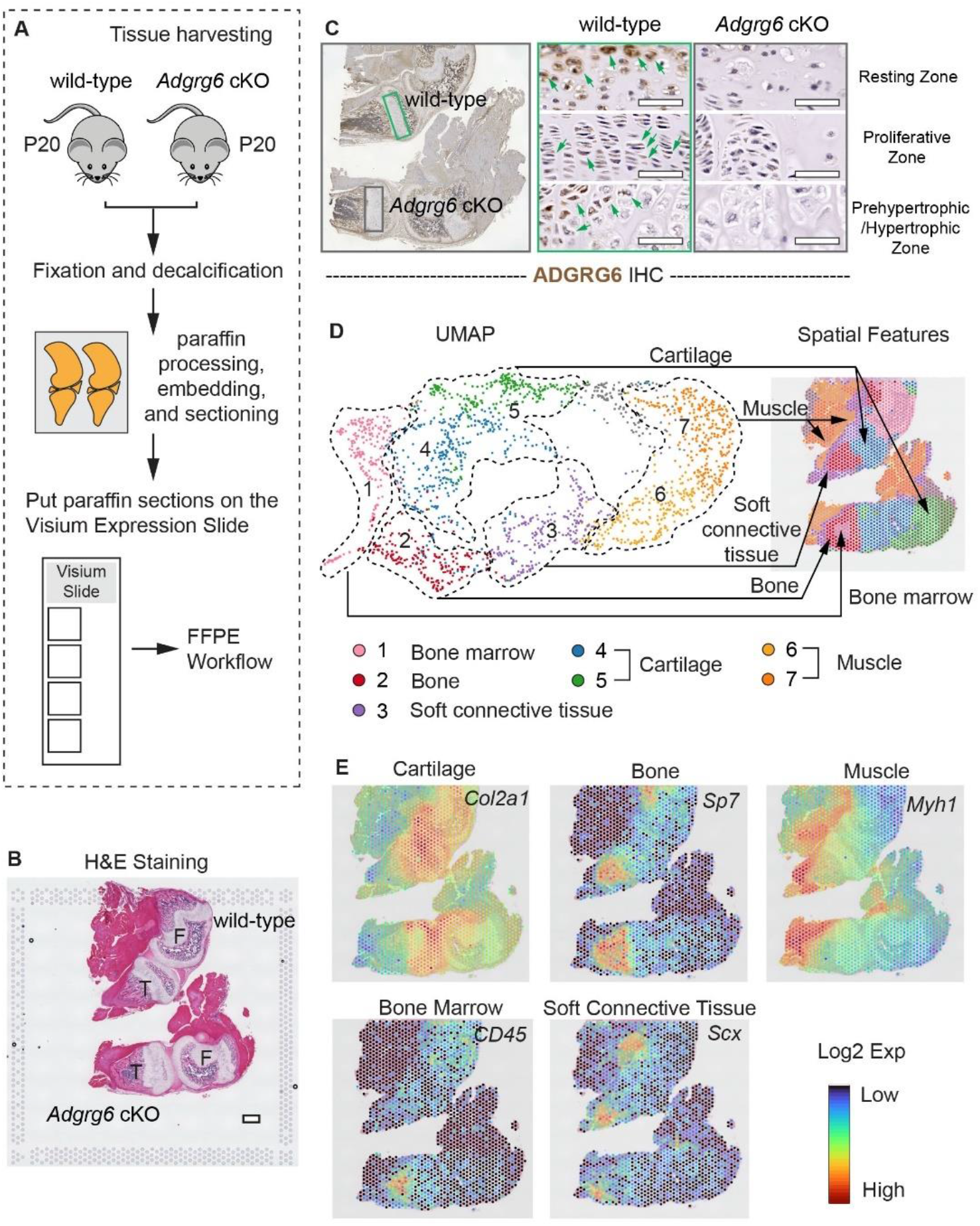
Establishment of a spatial transcriptomics workflow on the postnatal knee joint. **(A)** Schematic of the Visium FFPE workflow of the whole knee joint of P20 mice. **(B)** H&E staining of the wild-type control and *Adgrg6* cKO mice on the Visium gene expression slide. **(C)** IHC staining of ADGRG6 performed on the adjacent section for spatial transcriptomics analyses shows reduced ADGRG6 expression in all the zones of the *Adgrg6* mutant growth plate (green boxes) compared with the control (gray boxes). **(D)** UMAP and spatial feature plots of tissue populations that were identified from the control and mutant joints. 7 unsupervised clusters were identified as indicated. **(E)** Spatial expression plots of representative markers for cartilage (*Col2a1*), bone (*Sp7*), muscle (*Myh1*), bone marrow (CD45), and soft connective tissues (*Scx*). H&E: Hematoxylin and eosin. IHC: Immunohistochemistry; F: femur; T: tibia. Scale bar: 500µm in B; 50µm in C.

Each spot contained an average of 92,335 reads and 5,137 unique genes. Adjacent tissue sections were collected for IHC analyses to validate the ablation of *Adgrg6* (Fig. 2C). Non-supervised clustering of the spatial spots identified 7 tissue populations of the knee joints that were visualized on uniform manifold approximation (UMAP) plots (Fig. 2D and Supplementary Table 1), including bone marrow myeloid tissues (cluster 1), bone (cluster 2), soft connective tissues (cluster 3), cartilage (clusters 4, 5), and muscle (cluster 6: slow muscle, and cluster 7: fast muscle). Spatial expression plots show strong correlations between representative tissue markers and the histological structures, including cartilage (*Col2a1*), bone (*Sp7*), muscle (*Myh1*), bone marrow (*CD45/Ptprc*), and soft connective tissues (*Scx*) (Fig. 2E). Altogether, we successfully established a spatial transcriptomics workflow for mineralized mouse tissues.

### Spatial transcriptomic analyses reveal precocious chondrogenic-to-osteogenic conversion and increased expression of catabolic enzymes in the *Adgrg6* cKO mutant growth plates

Next, we conducted spatial transcriptomic analyses by analyzing the capture spots underlying the control and mutant growth plates (Fig. 3A). 256 differentially expressed genes were identified (Fig. 3B and Supplementary Table 2). Analyses of gene ontology (GO) enrichment for biological processes showed that ossification, skeletal system development, and extracellular matrix organization were predominantly affected (Fig. 3C and Supplementary Table 3). Detailed analyses of the top enriched, differentially expressed genes revealed that the expression of osteogenic genes (e.g., *Col1a1*, *Ibsp*, *Spp1*, *Sp7*, *Runx2, and Postn*) is significantly increased in the *Adgrg6* mutant growth plate (Fig. 3B, D, F and Supplementary Table 2). For instance, spatial expression analysis showed increased expression of *Spp1*, which encodes osteopontin, in the *Adgrg6* mutant growth plate (Fig. 3F). Osteopontin (SPP1) is an ECM glycoprotein highly expressed in osteoblasts and osteocytes ^31^, which can also be detected in some wild-type hypertrophic chondrocytes (red arrows, Fig. 3F’ left panel); however, IHC analysis showed that its expression is substantially increased throughout the *Adgrg6* mutant growth plates (red arrows, Fig. 3F’ right panel). These data suggest that the chondrocytes in the *Adgrg6* mutant growth plates precociously initiate the transition toward osteoblastogenesis, mimicking the phenotypes observed in the conditional *Sox9* knockout mice ^11^.

**Figure 3.**
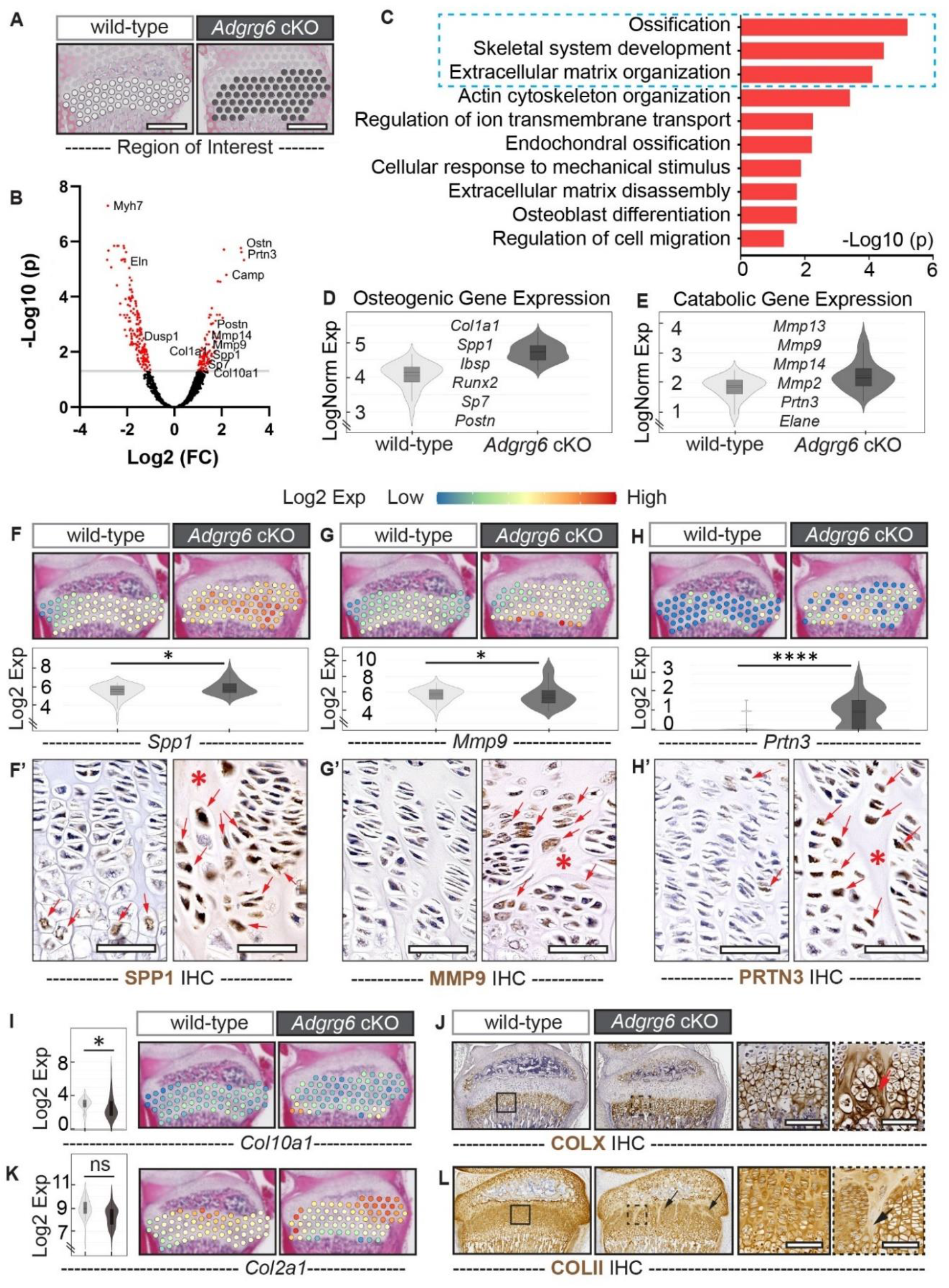
Spatial transcriptomic analyses reveal precocious chondrogenic-to-osteogenic conversion and increased catabolic gene expression in the *Adgrg6* cKO mutant growth plates. **(A)** Spatial capture spots underlying the wild-type (white spots) and *Adgrg6* cKO (gray spots) growth plates are selected for spatial transcriptomic analyses. **(B)** The volcano plot of differentially expressed genes (p<0.05, |Log2FC|>1). **(C)** GO enrichment analysis of the differentially expressed genes revealed altered biological processes (the top three are boxed in blue). **(D, E)** Violin plots of the Log normalized average expression (LogNorm Exp) of differentially expressed osteogenic and catabolic genes. (Osteogenic gene expression: wild-type, max: 4.669, q3: 4.295, mean: 4.144; *Adgrg6* cKO, max: 5.315, q3: 4.917, mean: 4.720. Catabolic gene expression: wild-type, max: 2.550, q3: 2.216, mean: 2.008; *Adgrg6* cKO, max: 3.783, q3: 2.595, mean: 2.365.) **(F-H’)** Spatial expression plots and violin plots of *Spp1*, *Mmp9*, and *Prtn3*. (*Spp1*: *p=0.0148; *Mmp9*: *p=0.0120; *Prtn3*: ****p<0.0001). IHC analyses of SPP1 (F’), MMP9 (G’), and PRTN3 (H’) show increased protein expression in mutant growth plates (red arrows). Acellular clefts are indicated with red asterisks. **(I, J)** Violin plots and spatial expression plots of *Col10a1* (I) and IHC analysis of COLX (J). (*Col10a1*: *p=0.0494.) The red arrow indicates the COLX-positive extracellular matrix. **(K, L)** Violin plots and spatial expression plots of *Col2a1* (K) and IHC analysis of COLII (L). (*Col2a1*: ns, p=1.0.) The reduced COLII signal around the acellular clefts is indicated with black arrows. (n=3 per genotype.) IHC: Immunohistochemistry. Scale bar: 500µm in A; 100µm in F’, G’, H’, J, and L.

We also observed increased expression of several catabolic enzymes (e.g., *Mmp13*, *Mmp9*, *Mmp14*, *Mmp2*, *Prtn3*, and *Elane*) in the *Adgrg6* mutant growth plate (Fig. 3E). IHC analyses confirmed increased expression of MMP9 and PRTN3 (proteinase 3) (Fig. 3G-H’). Some cells that highly express these catabolic enzymes are adjacent to the acellular clefts (red arrows and red asterisks, Fig. 3G’ and H’), suggesting that abnormal ECM remodeling may contribute to the disorganized growth plates in the *Adgrg6* mutants. GO enrichment analyses also revealed altered gene expression involved in cytoskeleton organization (*Actn3*, *Fscn1, Flnc*), glycolysis (*Pkm*, *Pgam2*, *Pgk1*), inflammatory response (*S100a8*, *S100a9, Camp*), ion transport (*Cacnb1*, *Kcnc1, Hvcn1*), and cell migration (*Macf1*, *Tnn*, *Plxnb2*) (Supplementary Table 3).

Since the *Adgrg6* cKO mice exhibit elongated hypertrophic zones (Figure 1I’), we next checked the expression of *Col10a1*. Spatial expression plots and IHC analyses showed increased *Col10a1* expression and expanded COLX staining in the mutant growth plates (Fig. 3I, J). No obvious change was observed with *Col2a1* expression by spatial expression analysis (Fig. 3K), but IHC staining revealed regionally reduced COLII signal in the mutant growth plates, especially around the acellular clefts (Fig. 3L). Altogether, our results suggest that ADGRG6 plays a crucial role in maintaining the chondrocyte fate of the postnatal growth plates. Loss of *Adgrg6* leads to a precocious chondrogenic-to-osteogenic conversion and ectopic upregulation of catabolic enzymes in the growth plate chondrocytes.

### Loss of *Adgrg6* results in impaired SOX9 expression and induced IHH signaling in growth plate chondrocytes

Next, we checked the expression of *Sox9*, one of the master regulators of cartilage maintenance and chondrocyte plasticity ^11, 32^. Spatial expression analysis showed no significant difference in *Sox9* expression between the control and mutant samples (Fig. 4A, F). However, IHC analysis revealed that SOX9 expression is reduced in the mutant growth plates in a region-specific manner. The percentage of SOX9 (+) cells is decreased in the resting zone of the mutant growth plates (Fig. 4H) but is not significantly altered in the proliferative zone (Fig. 4H), which is largely due to the presence of SOX9 (+) cells adjacent to the acellular clefts (green arrows, Fig. 4G mutant panel). About 40% of cells in the wild-type hypertrophic growth plates are SOX9 (+), but this percentage is reduced to 15% in the mutants (Fig. 4G, H). Taken together, we show that loss of *Adgrg6* leads to decreased SOX9 expression in the resting and hypertrophic chondrocytes of the growth plates.

**Figure 4.**
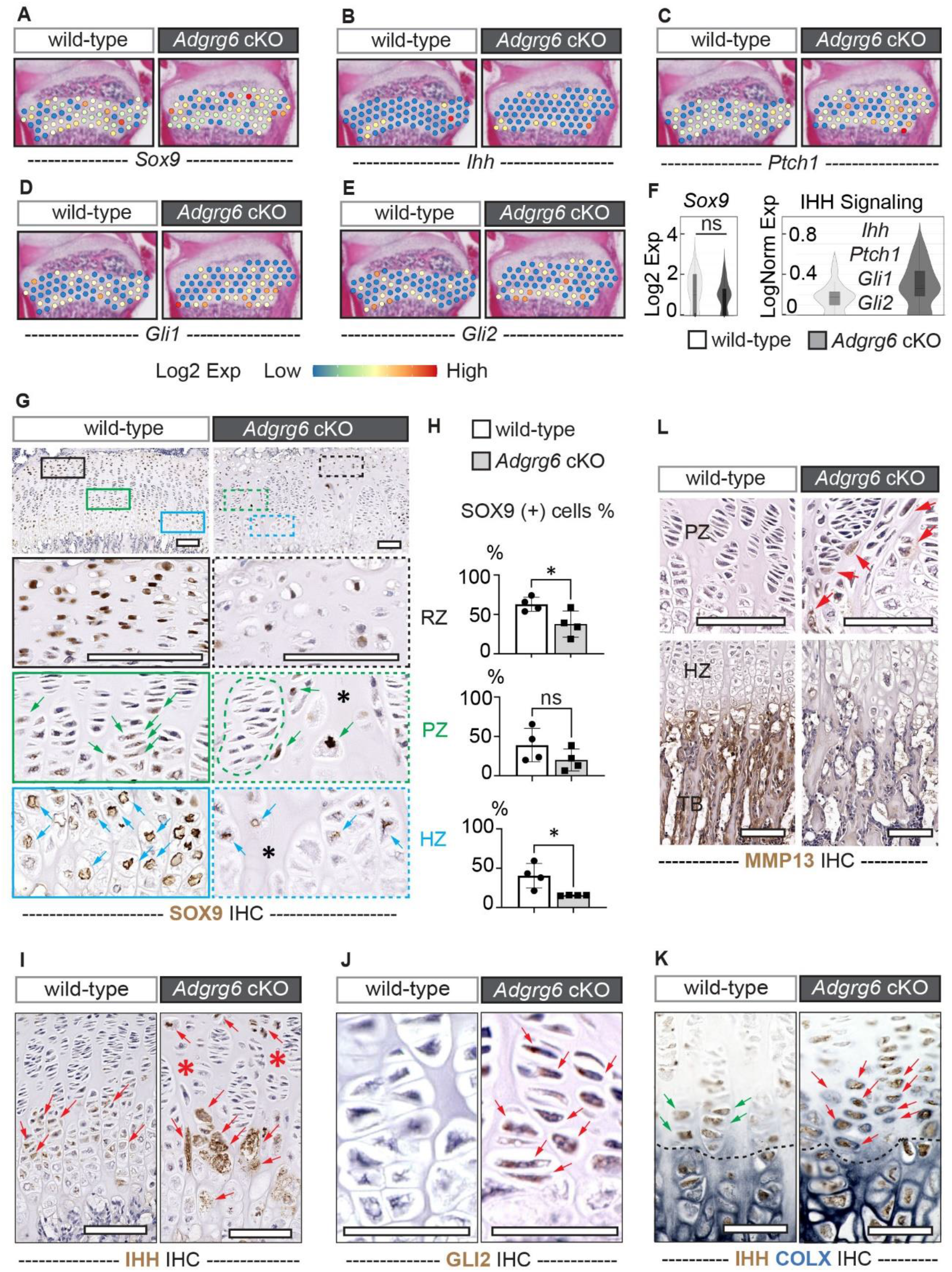
Loss of *Adgrg6* results in impaired SOX9 expression and induced IHH signaling in growth plate chondrocytes. **(A-F)** Spatial expression plots of *Sox9* (A) and IHH signaling molecules *Ihh* (B), *Ptch1* (C), *Gli1* (D), and *Gli2* (E) on P20 growth plates. The violin plots of *Sox9* and the Log normalized average expression (LogNorm Exp) of IHH signaling are shown in F. (*Sox9*: ns, p=1.0. IHH signaling: wild-type, max: 0.539, q3: 0.229, mean: 0.182; *Adgrg6* cKO, max: 0.743, q3: 4.434, mean: 0.292.) **(G, H)** IHC analyses of SOX9 revealed reduced expression in the resting and hypertrophic zones (black and blue boxes) but not the proliferative zone (green boxes) of the mutant growth plates at P20. SOX9 (+) cells are indicated with green and blue arrows. Cells formed into clusters are circled with a green dashed line. Acellular clefts are indicated with black asterisks. The percentage of SOX9 (+) cells in each zone is quantified in H. (n=4 per genotype. *p=0.0374 in RZ; ns, p=0.1880 in PZ; *p=0.0189 in HZ.) Bars are plotted by means ± SD, two-tailed *t*-test. **(I-K)** IHC analyses of IHH (I), GLI2 (J), and dual color IHC of IHH (brown signal) and COLX (dark blue signal) (K) are shown in control and mutant growth plates at P20. IHH (+) cells and acellular clefts are indicated with red arrows and red asterisks in I. GLI2 (+) cells are indicated with red arrows in J. COLX (+) cells in the mutant prehypertrophic growth plate are indicated with red arrows in K; COLX (-) cells in the control prehypertrophic growth plate are indicated with green arrows in K. The black dashed lines in K indicate the boundary between the prehypertrophic and hypertrophic zones. (n=4 per genotype.) **(L)** IHC analysis of MMP13 in the proliferative and terminal hypertrophic growth plates of the control and mutant mice at P20. MMP13 (+) cells are indicated with red arrows. (n=4 per genotype.) IHC: Immunohistochemistry. RZ: resting zone; PZ: proliferative zone; HZ: hypertrophic zone; TB: trabecular bone. Scale bar: 100µm in G, I, L, and M; 50µm in J, K.

The elongated prehypertrophic/hypertrophic growth plates in the *Adgrg6* cKO mice partially recapitulate the phenotypes of the conditional *Pth1r* knockout mice, which exhibit an elongated hypertrophic zone that expresses *Ihh* before the premature closure of the growth plates ^22, 23^. Spatial expression analyses revealed that the IHH signaling molecules (e.g. *Ihh*, *Ptch1*, *Gli1*, and *Gli2*) are upregulated in the *Adgrg6* mutant growth plate (Fig. 4B-F). IHC staining showed that IHH expression is more diffused in the mutant growth plates, ranging from the prehypertrophic/hypertrophic chondrocytes to the proliferative chondrocytes (red arrows, Fig. 4I right panel), in contrast to the more restricted expression in the prehypertrophic and early hypertrophic cells of the control growth plates (red arrows, Fig. 4I left panel). The activation of the IHH signaling pathway was confirmed by increased GLI2 IHC staining in the mutant growth plates (Fig. 4J). To better understand how the expanded IHH signaling contributes to hypertrophic differentiation, we performed dual-color IHC analyses of IHH (brown signal, Fig. 4K) and COLX (dark blue signal, Fig. 4K). In control mice, the expression of COLX is more restricted to the enlarged hypertrophic cells (cells below the black dashed line, Fig. 4K left panel), and the IHH positive cells in the prehypertrophic region are largely COLX (-) (green arrows, Fig. 4K left panel). However, in the mutants, we consistently observed increased COLX expression in the pericellular matrix of the IHH-positive prehypertrophic cells (red arrows, Fig. 4K right panel). These observations suggest that the ectopic IHH signaling in the *Adgrg6* mutant mice may promote premature hypertrophic differentiation.

We next examined the expression of MMP13 (Matrix Metallopeptidase 13) which resorbs hypertrophic cartilage during endochondral ossification ^1^. Interestingly, MMP13 expression is also altered in a region-specific manner upon loss of *Adgrg6*. We observed increased, ectopic MMP13 expression in the mutant proliferative/prehypertrophic growth plates (red arrows, Fig. 4L), which is consistent with our spatial expression analyses (Fig. 3E). However, MMP13 expression is reduced in the terminal hypertrophic growth plate and the primary spongiosa/trabecular bone of the mutants (Fig. 4L), suggesting an impaired remodeling of the terminal hypertrophic chondrocytes. These data indicate that the accumulation of hypertrophic cells in the *Adgrg6* cKO mice may be due to induced prehypertrophic/hypertrophic differentiation caused by increased IHH signaling, and impaired terminal hypertrophic maturation caused by reduced MMP13 expression. Taken together, these data show that loss of *Adgrg6* results in impaired SOX9 expression and induced IHH signaling in growth plate chondrocytes, which may contribute to the precocious chondrogenic-to-osteogenic conversion and dysregulated hypertrophic differentiation in the *Adgrg6* mutant growth plates.

### Spatial analyses highlight altered cell populations and changes in cell-cell crosstalk in the *Adgrg6* cKO growth plates

To better understand the regulatory mechanisms of ADGRG6 in different growth plate cell populations, we sub-clustered the capture spots underlying the control and mutant growth plates. UMAP of gene expression profiles identified three cell populations (clusters 0-2) as shown on the tissue sections (Fig. 5A, B). One limitation of this analysis is that the spatial transcriptomics is not at single-cell resolution, where one capture spot covers multiple cells and may span the junction of different zones of the growth plate (Supplementary Fig. 2A, B, C). Therefore, the cell populations identified based on the capture spots may contain a mixture of cell types. Despite these caveats, we decided to see if unbiased clustering could reveal any alterations in molecular signaling, which could be verified by orthogonal approaches downstream.

**Figure 5.**
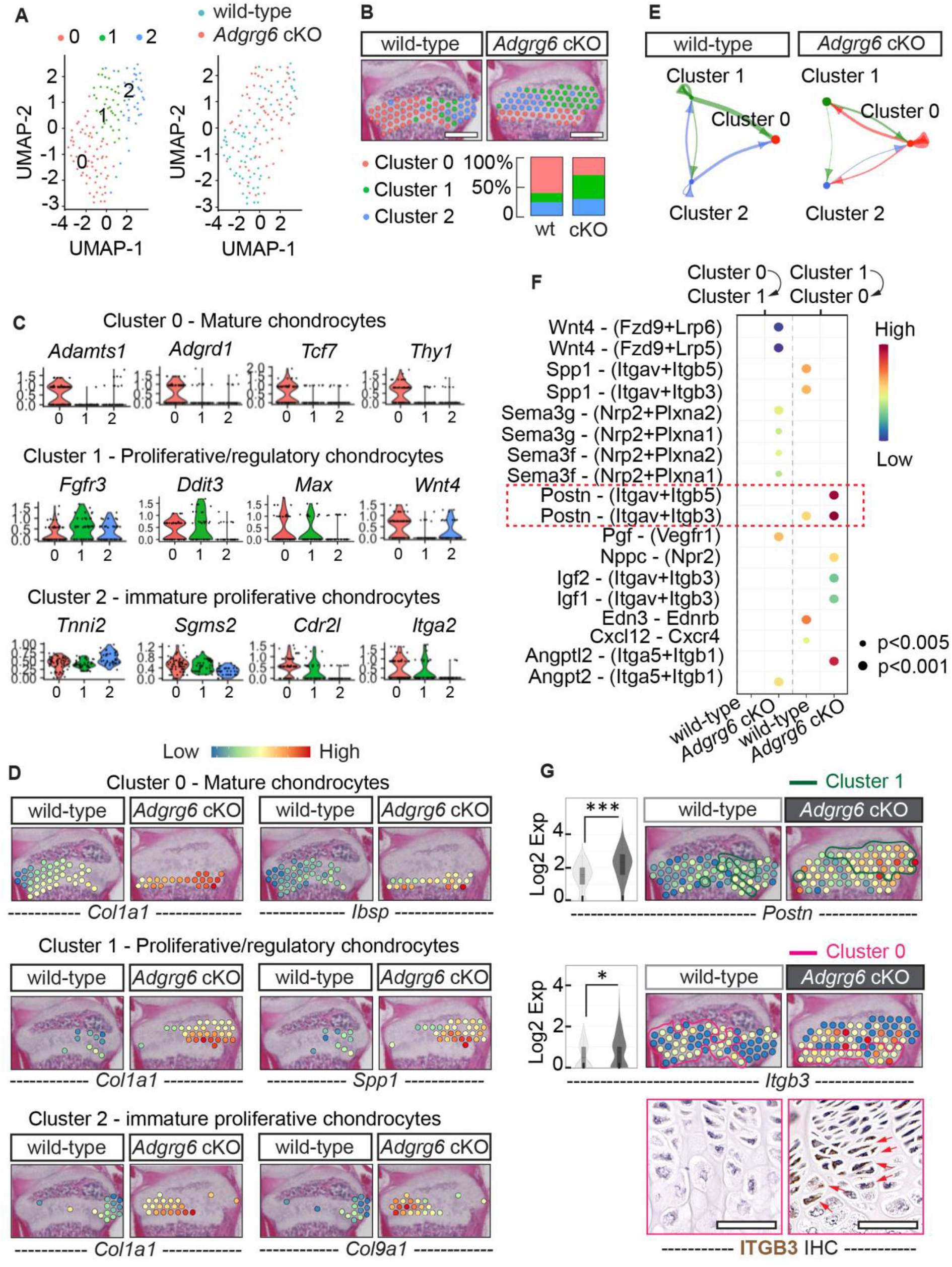
Spatial analyses highlight altered cell populations and changes in cell-cell crosstalk in the *Adgrg6* cKO growth plates. **(A)** UMAP plots of cell populations from the wild-type and the *Adgrg6* cKO mutant growth plates. Three clusters (clusters 0, 1, 2) were identified. **(B)** The spatial distribution and the percentage of each cluster are shown. **(C)** Violin plots show the representative markers of each cluster. **(D)** Several top differentially expressed genes in each cluster are shown with spatial expression plots. **(E)** The communications of significant ligand-receptor interactions among the three clusters are shown. The dots’ size and the edges’ width are proportional to the number of capture spots in each cluster and the number of ligand-receptor pairs, respectively. **(F)** The dot plot shows all the significant ligand-receptor pairs that are involved in the crosstalk between cluster 0 and cluster 1. The dot color and size represent the calculated communication probability and p-values. p-values are computed from the one-side permutation test. **(G)** Violin plots and spatial expression plots of the top enriched ligand-receptor pair, *Postn*-*Itgb3*, and IHC analysis of ITGB3 are shown. Capture spots in cluster 1 and cluster 0 are circled with green lines and magenta lines, respectively. (*Postn*: ***p= 0.0009; *Itgb3*: *p= 0.0235.) ITGB3 (+) cells in the mutant growth plates are indicated with red arrows. (n=4 per genotype.) IHC: Immunohistochemistry. Scale bar: 500µm in B; 50µm in G.

Analysis of conserved marker in each cluster suggests that cluster 0 defines a group of mature chondrocytes, as it expresses the ECM degrading enzyme (*Adamts1*) ^33^ and the bone-enriched GPCR (*Adgrd1*/*Gpr133*) which is associated with bone mineral density ^34^. It also highly expresses the Wnt signaling-related gene *Tcf7* which is involved in osteogenesis ^35^ and the skeletal progenitor marker *Thy1/CD90* ^36^. The percentage of cluster 0 spots is 62% in the control growth plate, but it is decreased to 15% in the mutant growth plate with distribution more toward the hypertrophic zone (Fig. 5B). Cluster 1 spots are defined as proliferative/regulatory chondrocytes, which expresses proliferative cell marker (*Fgfr3*) ^37^ and genes that are highly expressed in the regulatory chondrocytes in human cartilage tissues (*Ddit3*, *Max*) ^38^. Cluster 1 spots are low in the expression of the Wnt signaling molecule (*Wnt4*) that is involved in hypertrophic differentiation ^39^, suggesting that these chondrocytes are less mature. Cluster 1 spots compose 15% of the spots in the control growth plate that are sparsely located in the resting and proliferative zones; however, the percentage of cluster 1 cells is increased to 41% in the *Adgrg6* cKO mutants growth plate (Fig. 5B). Cluster 2 defines a group of immature proliferative chondrocytes that highly express *Tnni2*, a gene member of the troponin I gene family that functions as an angiogenesis inhibitor in cartilage ^40^. *Tnni2* expression is highly enriched in the resting and immature proliferative chondrocytes but is absent in mature proliferative chondrocytes in mice ^41^. Cluster 2 spots have low expression of genes that are prominently expressed in bone (*Sgms2*, *Spon1*) ^42, 43^ or skeletal progenitors correlated with ossification-commitment (*Itga2*) ^44^. 23% and 29% of the spots are classified as cluster 2 spots in control and mutant growth plates, respectively (Fig. 5B). Altogether, the altered distribution of cell clusters in *Adgrg6* mutants shows more enrichment of chondrocytes at a less differentiated or dedifferentiated stage (clusters 1 and 2) and a reduction of cells at the mature chondrocyte stage (cluster 0).

To better understand the nature of these altered cell types, we analyzed the differentially expressed genes in each cluster (Supplementary Table 5). We found that the osteogenic marker *Col1a1* is upregulated in all three clusters of the *Adgrg6* mutant growth plate (Fig. 5D). Additional known osteogenic markers, including *Ibsp*, *Spp1*, *Col1a1*, *Dmp1,* and *Bmp1* are upregulated in cluster 0 spots (mature chondrocytes), while *Serpinh1*, *Spp1*, *Ibsp*, *Ostn*, and *Postn* are upregulated in cluster 1 spots (proliferative/regulatory chondrocytes) (Fig. 5D and Supplementary Table 5). The expression of *Col9a1*, which is more enriched in fetal cartilage compared with adult cartilage ^45^, is increased in cluster 2 (immature proliferative chondrocytes) of the mutant growth plate (Fig. 5D), suggesting that these cells are either delayed in differentiation or have been dedifferentiated into less mature chondrocytes. Taken together, these data confirm that ADGRG6 is required for regulating proper differentiation and maintaining chondrocyte identity in all the zones of the postnatal growth plates.

Next, we performed CellChat ^46^ analyses to highlight candidate signaling pathways (receptor-ligand pairs) that are co-regulated, in comparison of each cluster between control and mutant growth plates. We found that in the control growth plate, cluster 0 cells (mature chondrocytes) are mainly receiver cells, acquiring signaling molecules (e.g., ligands) sent by cluster 1 and cluster 2 cells (proliferative/regulatory, or immature proliferative cells) (Fig. 5E). However, in the mutant growth plates, cluster 0 cells act as both senders and receivers, exhibiting increased interfered interactions with themselves and with cells from other clusters, and show a stronger communication with cluster 1 cells (Fig. 5E, Supplementary Fig. 3A). Detailed analyses of ligand-receptor pairs involved in the crosstalk between cluster 0 and cluster 1 predicted that the *Postn*-(*Itgav*+*Itgb5* or *Itgav*+*Itgb3*) communication is one of the most significant signaling pathways from cluster 1 to cluster 0 in the *Adgrg6* cKO mice (red box, Fig. 5F, Supplementary Fig.3C). This pattern is confirmed by the spatial expression plots and the IHC staining of ITGB3 (integrin subunit beta 3) (Fig. 5G). POSTN (periostin) is preferentially expressed in periosteal osteoblasts and osteocytes in response to PTH and can directly inhibit *Sost* expression through its integrin αVβ3 receptor (*Itgav*+*Itgb3*) ^47^. This increased communication between POSTN and integrin receptors was also observed between cluster 0 and cluster 2 (Supplementary Fig. 3C). Thus, the ectopic activation of key osteogenic receptor-ligand pairs, *Postn*-(*Itgav*+*Itgb5* or *Itgav*+*Itgb3*), suggests a molecular mechanism promoting the abnormal osteoblastogenesis in the *Adgrg6* mutant growth plates. In addition, we observed evidence of increased communication of Wnt [*Wnt4*-(*Fzd9*+*Lrp5* or *Fzd9+Lrp6*)] and Semaphorin signaling [*Sema3g*/*Sema3f*-(*Nrp2*+*Plxna1*/*Plxna2*)] in the mutant cell clusters (Fig. 5F, Supplementary Fig. 3B, C), suggesting that the hypertrophic differentiation and cytoskeletal/adhesive machinery^48, 49^ of the mutant growth plates might also be affected.

### ADGRG6 is required for chondrocyte survival and the proper formation of the resting zone

GO enrichment analyses also revealed altered processes of “protein kinase A signaling” and “negative regulation of cell proliferation” (Supplementary Table 3). In agreement, we observed attenuated Gs/cAMP/PKA/CREB signaling, supported by reduced expression of phosphorylated CREB (pCREB) (Supplementary Fig. 4A, B); and decreased cell proliferation, evidenced by decreased expression of Ki67, in the *Adgrg6* mutant growth plates at P20 (Supplementary Fig. 4C, D). As Gs and Gq/11 signaling play an important role in maintaining cell survival within the resting zone ^23, 24^, we next checked if cell death was altered in *Adgrg6* cKO growth plates. TUNEL staining revealed increased cell death in all the zones of the mutant growth plates at P20 (Supplementary Fig. 4E, F), indicating that ADGRG6 is required for maintaining growth plate chondrocyte survival.

To better understand how ADGRG6 regulates the establishment and maintenance of the resting zone chondrocytes, we examined the growth plates of the control and *Adgrg6* cKO mice at P14, when the secondary ossification center and the resting zone are fully developing in the wild-type mice. Histological analysis revealed that the *Adgrg6* cKO mice exhibit delayed formation of the secondary ossification center at P14 (Fig. 6A-B) and reduced cellularity in the resting and proliferative growth plates (Fig. 6A-B’, E). Abnormal formation of cell clusters and acellular clefts were also observed in the mutants (Fig. 6B’). TUNEL staining revealed increased cell death in all the zones of the mutant growth plates at P14 (Fig. 6C-D’, F), especially in the cells that formed into clusters within the resting zone (yellow circles, Fig. 6D’). The increased cell death is associated with a significantly reduced number of ADGRG6 and SOX9-positive cells within the resting zone of the mutant growth plates (Fig. 6G-J).

**Figure 6.**
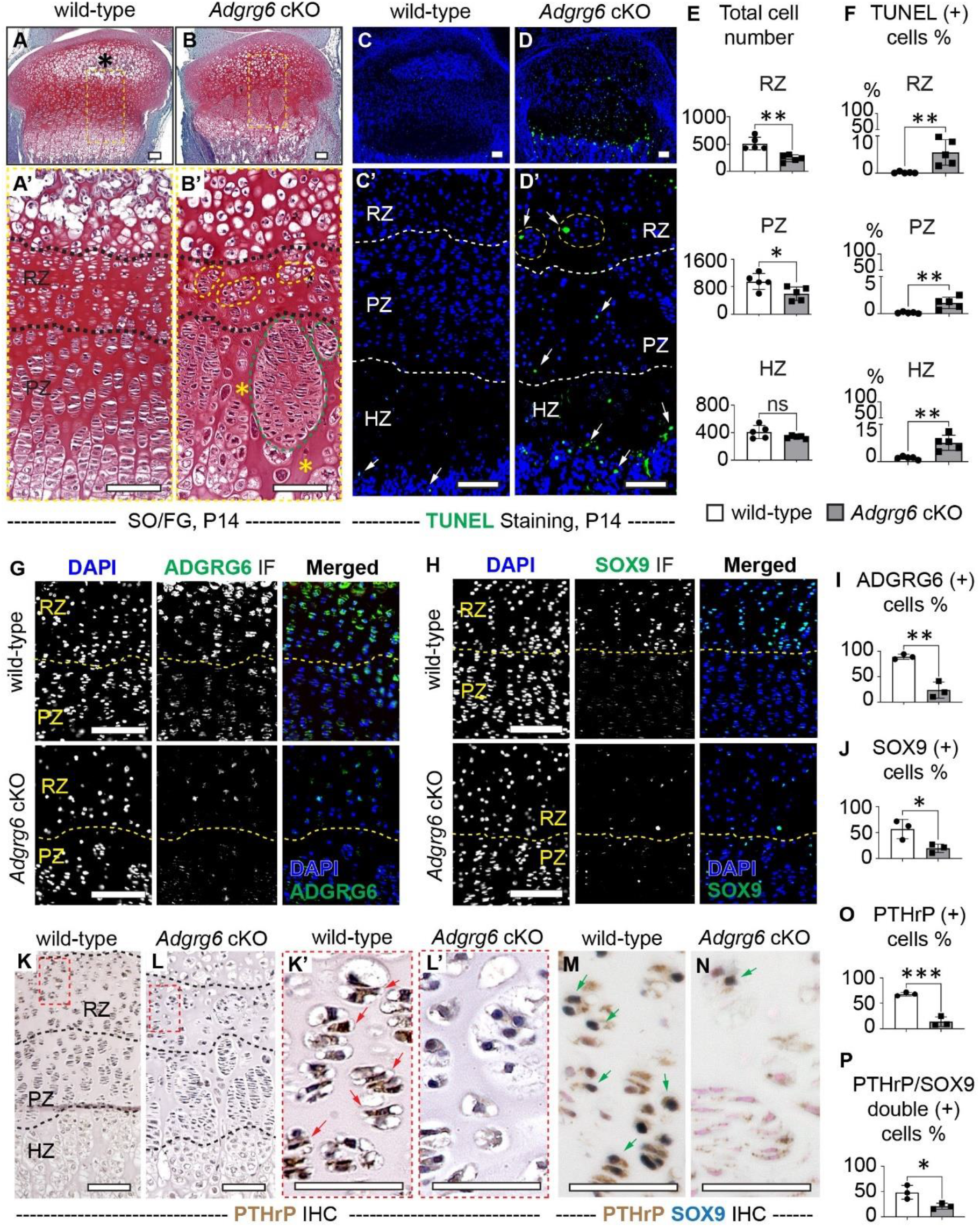
ADGRG6 is required for chondrocyte survival and the proper formation of the resting zone. **(A-B’)** SO/FG staining of the wild-type and *Adgrg6* ckO growth plates at P14. Normally formed secondary ossification center in the control is indicated with a black asterisk. Cells formed into clusters in the mutant resting and proliferative zones are circled with yellow and green dashed lines. Acellular clefts are indicated with yellow asterisks. (n=5 per genotype.) **(C-D’)** TUNEL staining of control and mutant growth plates at P14. TUNEL (+) cells are indicated with white arrows. Cells formed into clusters in the resting zone are circled with yellow dashed lines in D’. (n=5 per genotype.) **(E-F)** The total cell number and the percentage of TUNEL (+) cells in each zone are quantified. (n=5 per genotype. E: **p= 0.0019 in RZ; *p= 0.0302 in RZ; ns, p=0.1267 in HZ. F: **p= 0.0081 in RZ; **p= 0.0050 in RZ; **p= 0.0022 in HZ.) (**G-J**) IF analyses of ADGRG6 (G) and SOX9 (H) in control and mutant growth plates at P14. The grayscale channels and the merged DAPI and ADGRG6/SOX9 IF channels are shown. The percentage of ADGRG6 (+) and SOX9 (+) cells in the resting zone are quantified in I and J. (n=5 per genotype. I: **p= 0.0024; J: *p= 0.0343). (**K-P**) IHC analysis of PTHrP (K-L’) and dual color IHC analysis of PTHrP (brown signal) and SOX9 (dark blue signal) are shown in control and mutant growth plates (M, N, counterstained with nuclear fast red) at P14. PTHrP (+) cells are indicated with red arrows in K’, and PTHrP/SOX9 double (+) cells are indicated with green arrows in M and N. The percentage of PTHrP (+) cells and PTHrP/SOX9 double (+) cells in the resting zone is quantified in O and P. (n=3 per genotype. O: ***p= 0.0007; P: *p= 0.0334.) Bars are plotted by means ± SD, two-tailed *t*-test. SO/FG: Safranin O/Fast Green. IHC: Immunohistochemistry. IF: immunofluorescence. RZ: resting zone; PZ: proliferative zone; HZ: hypertrophic zone. Scale bar: 100µm in A-D’, G, H, K, and L; 50µm in K’, L’, M, and N.

Next, we checked the expression of PTHrP, as the formation of the resting zone is closely associated with a group of *Pthrp* (+) chondrocytes in the postnatal growth plates ^4^. IHC analysis shows that at P14, PTHrP is highly expressed in the resting and proliferative chondrocytes of the control growth plates (Fig. 6K, K’); however, its expression is substantially reduced in the mutant growth plates (Fig. 6L, L’, O). The reduced PTHrP expression can also be observed in P20 *Adgrg6* cKO mice (Supplementary Fig. 5A). Dual color IHC analysis revealed that the SOX9 (+) cells (dark blue signal, Fig. 6M, N) are colocalized with the PTHrP (+) cells (brown signal, Fig. 6M, N) in the resting zone at P14, and the number of these SOX9/PTHrP double-positive cells is significantly reduced in the resting zone of the *Adgrg6* cKO mice (Fig. 6M, N, P, Supplementary Fig. 5B). The *Adgrg6* cKO mice also exhibit expanded expression of COLX and IHH in the prehypertrophic/hypertrophic and proliferative growth plates at P14 (Supplementary Fig. 5C, D). Taken together, our data suggest that ADGRG6 maintains chondrocyte survival and regulates resting zone formation and maintenance by controlling SOX9 and PTHrP expression.

### ADGRG6 is essential for the homeostasis of the postnatal growth plate

To better understand the role of *Adgrg6* specifically in postnatal growth plates, we generated mice carrying the *Adgrg6^f/f^* and the *AcanCre^ERT^*^2^ (Tamoxifen-inducible chondrocyte-specific Cre) alleles ^50^ (*AcanCre^ERT^*^2^*; Adgrg6^f/f^*, called *Adgrg6* cKO^TM^ hereafter). We treated control and *Adgrg6* cKO^TM^ mutant mice with Tamoxifen from P14 to P18 to delete *Adgrg6* expression after the formation of the secondary ossification center, and harvest knee joints 6 days post recombination at P20. Histological analysis showed a decreased length of the growth plates in the *Adgrg6* cKO^TM^ mice (Fig. 7A-B’), associated with reduced cellularity and decreased number of cell layers in the resting and proliferative zones (Fig. 7C-E). The efficient ablation of *Adgrg6* is demonstrated by reduced expression of ADGRG6 and pCREB in the *Adgrg6* cKO^TM^ mutant growth plates (Fig. 7F, J, K, Supplementary Fig. 6A, B). TUNEL staining showed increased cell death in the resting zone but not the proliferative or hypertrophic zones of the *Adgrg6* cKO^TM^ growth plates (Fig. 7G-I), suggesting that the primary function of ADGRG6 in postnatal growth plates is to regulate resting chondrocyte survival and homeostasis. In addition, alterations in gene expression observed in P20 *Adgrg6* cKO mice were rapidly recapitulated in *Adgrg6* cKO^TM^ mice at P20, including decreased expression of SOX9, more diffuse expression of IHH, and increased expression of osteogenic marker SPP1 and catabolic enzyme MMP9 (Fig. 7L-O, Supplementary Fig. 6C-F). Altogether, these data show that ADGRG6 plays an essential role in maintaining homeostasis of the postnatal growth plate chondrocytes.

**Figure 7.**
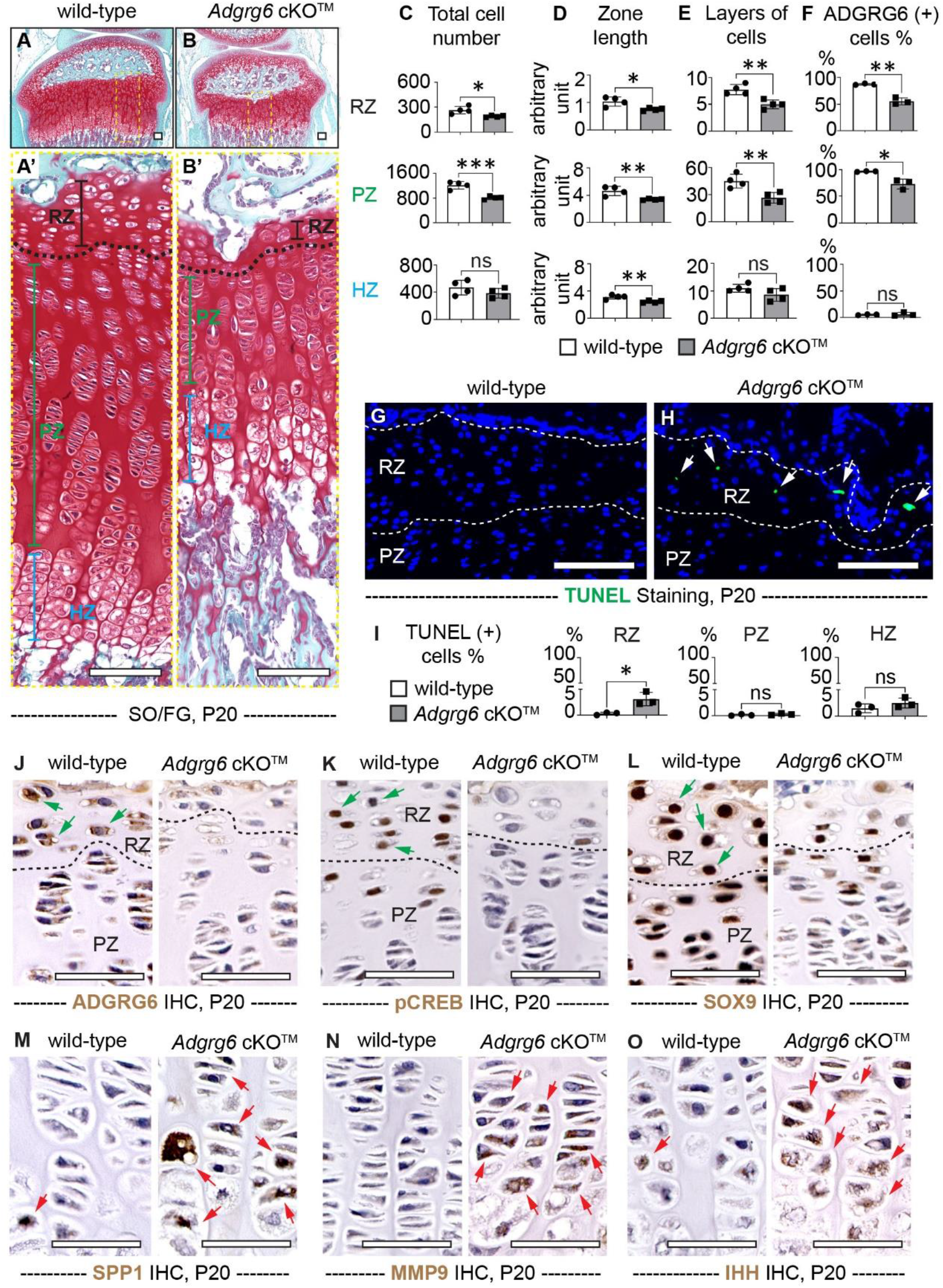
ADGRG6 is essential for the homeostasis of the postnatal growth plate. **(A-B’)** SO/FG staining of the wild-type and *Adgrg6* ckO^TM^ growth plates at P20. *Adgrg6* ablation with *AcanCre^ERT^*^2^ was induced with Tamoxifen induction from P14 to P18. The length of the resting zone, proliferative zone, and hypertrophic zone is indicated with a black, green, and blue segment, respectively. **(C-F)** The total cell number, zone length, and layers of cells in each zone of the growth plates are quantified in C-E. (n=4 per genotype. C: *p= 0.0184 in RZ; ***p= 0.008 in PZ; ns, p= 0.2409 in HZ. D: *p= 0.0169 in RZ; **p= 0.0099 in PZ; **p= 0.0070 in HZ. E: **p= 0.0047 in RZ; **p= 0.0086 in PZ; ns, p= 0.1210 in HZ.) The percentage of ADGRG6 (+) cells in each zone of the growth plate is quantified in F. (n=3 per genotype. F: **p= 0.0011 in RZ; *p= 0.0114 in PZ; ns, p= 0.7329 in HZ.) **(G-I)** TUNEL staining of control and mutant growth plates at P20 (G, H). TUNEL (+) cells are indicated with white arrows. The percentage of TUNEL (+) cells in each zone is quantified in I. (n=3 per genotype. I: *p= 0.0214 in RZ; ns, p= 0.3231 in PZ; ns, p= 0.2375 in HZ.) (**J-O**) IHC analyses show reduced expression of ADGRG6 (J), pCREB (K), and SOX9 (L), but increased expression of SPP1 (M), MMP9 (N), and IHH (O) in the mutant growth plates at P20. (n=3 per genotype.) Bars are plotted by means ± SD, two-tailed *t*-test. SO/FG: Safranin O/Fast Green. IHC: Immunohistochemistry. RZ: resting zone; PZ: proliferative zone; HZ: hypertrophic zone. Scale bar: 100µm in A-B’; 50µm in J-O.

## Discussion

Here we show that ADGRG6 is dispensable for embryonic limb development but is required for postnatal growth plate homeostasis. Under physiological conditions, ADGRG6 can promote Gs/CREB signaling and maintain normal expression of SOX9 and PTHrP to safeguard the identity of the growth plate chondrocytes, ensure proper formation of the resting zone, and suppress premature hypertrophic differentiation via interaction with IHH (Fig. 8A). Ablation of *Adgrg6* results in attenuated Gs/CREB signaling and reduced SOX9 expression, which may contribute to the precocious chondrogenic-to-osteogenic conversion and increased cell death. Loss of *Adgrg6* also results in reduced PTHrP expression and ectopic IHH signaling, which may further dysregulate the homeostasis of the resting chondrocytes and lead to increased flux towards prehypertrophic/hypertrophic differentiation. Finally, *Adgrg6* ablation leads to ectopic expression of several catabolic enzymes (e.g., MMP13) which may contribute to the ECM disorganization and dysregulated terminal hypertrophic differentiation (Fig. 8B). Together, we uncovered that ADGRG6 is required to maintain normal differentiation and survival of the postnatal chondrocytes. Together these dysregulated signaling pathways and biological processes disrupt the homeostasis of the postnatal growth plate.

**Figure 8.**
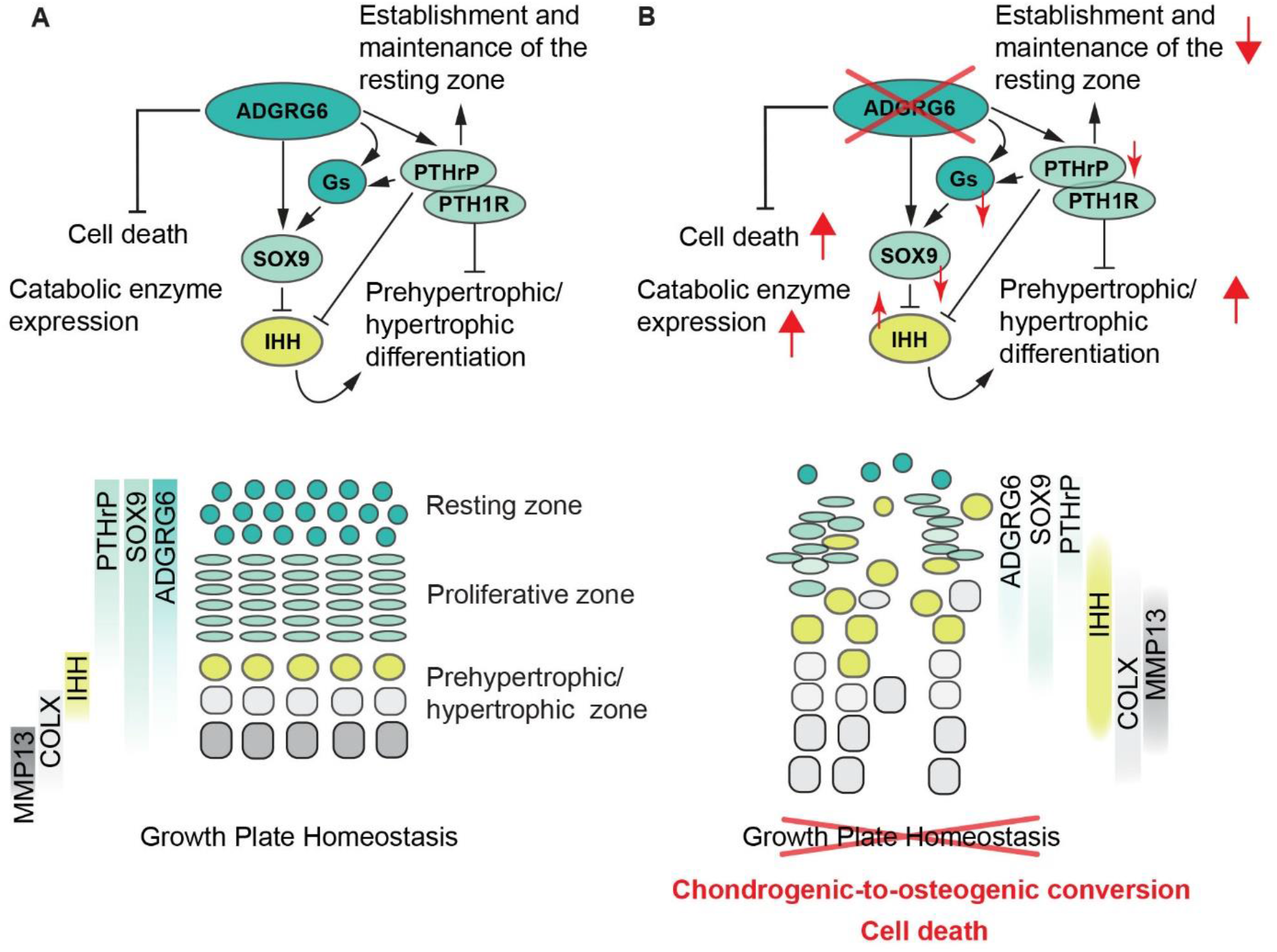
Proposed mechanisms of how ADGRG6 regulates postnatal growth plate homeostasis. **(A)** Under physiological conditions, ADGRG6 can signal through Gs signaling and maintain normal expression of SOX9 and PTHrP, while suppressing the IHH signaling. These signaling pathways safeguard chondrocyte identification and ensure proper formation of the resting zone; while suppressing premature hypertrophic differentiation, abnormal cell death, and excessive catabolic enzyme expression. **(B)** Upon the ablation of *Adgrg6*, Gs signaling is attenuated and the expression of SOX9 and PTHrP is reduced, while the expression of IHH and COLX is expanded associated with ectopic upregulation of catabolic enzymes including altered MMP13 expression (upregulated in the proliferative/prehypertrophic chondrocytes but reduced in the terminal hypertrophic chondrocytes). These changes lead to precocious chondrogenic-to-osteogenic conversion, increased cell death, promoted prehypertrophic/hypertrophic differentiation but impaired terminal hypertrophic differentiation, and overall disorganized growth plate structure.

Our study suggests that ADGRG6 interacts with both SOX9 and PTHrP/PTH1R/IHH signaling in postnatal chondrocytes. It is known that postnatal ablation of *Sox9* (*Sox9* cKO) ^11, 51^ or *Pth1r* (*Pth1r* cKO) ^22, 23^ leads to premature growth plate closure. Interestingly, though the *Adgrg6* cKO mice do not display a complete growth plate closure, they exhibit a similar chondrogenic-to-osteogenic conversion phenotype as observed in the *Sox9* cKO mice ^11^, and a premature hypertrophic differentiation phenotype as observed in the *Pth1r* cKO mice ^22^. Given that ADGRG6 is highly expressed in the resting and proliferative chondrocytes and exhibits an expression gradient opposing to PTH1R, our study suggests that ADGRG6 may play a cell-autonomous role in maintaining the homeostasis of the resting and proliferative chondrocytes by regulating SOX9 and PTHrP, but regulates hypertrophic chondrocytes via modulating other signaling molecules, such as IHH, in a non-cell-autonomous manner.

Haseeb and colleagues found that postnatal ablation of *Sox9* promotes growth plate chondrocytes in all the stages to quickly dedifferentiate and then convert into osteoblasts ^11^. During this process, cells tend to skip the prehypertrophic/hypertrophic differentiation as the expression of *Ihh* and *Col10a1* is quickly diminished. We observed a similar chondrogenic-to-osteogenic conversion phenotype in the resting and proliferative zones of our *Adgrg6* cKO mice, evidenced by increased expression of osteogenic markers (e.g., *Runx2*, *Sp7*, *Col1a1*, *Postn*, *Mmp13*, and *Ibsp*) that are all upregulated in the *Sox9* cKO mice ^11^. Our study also predicts that the induced POSTN/integrin receptor interaction may contribute to the formation of these precocious osteogenic cell types. Moreover, postnatal loss of *Sox9* leads to reduced *Adgrg6* expression in mouse IVDs ^51^, and conversely, *Adgrg6* ablation decreases SOX9 expression in the cartilaginous endplate and vertebral growth plate of the spine ^25^, suggesting that SOX9/ADGRG6 may interact in a positive feedback loop. In addition, CREB can bind to the human *SOX9* promoter ^52^ and the PTHrP/PKA signaling pathway can stimulate the phosphorylation of SOX9 and enhance its DNA-binding and transcriptional activity ^53, 54, 55^. Together, our results suggest that ADGRG6 can positively regulate SOX9 expression in growth plate chondrocytes, potentially through the Gs/PKA/CREB signaling pathway.

It is worth mentioning that loss of *Adgrg6* does not lead to complete ablation of SOX9 expression. Thus, our data may suggest that resting and proliferative cells are more sensitive to SOX9 dosage as incomplete loss of SOX9 expression is sufficient to induce chondrogenic-to-osteogenic conversion. In contrast, prehypertrophic/hypertrophic chondrocytes appear to be more tolerant to SOX9 reduction, so the residual SOX9 expression upon loss of *Adgrg6* is sufficient to initiate hypertrophic differentiation. This is supported by the altered cluster distribution in the *Adgrg6* cKO growth plates, where more cells in the resting and proliferative zones are calcified as less differentiated/dedifferentiated cells, but cells in the prehypertrophic/hypertrophic zones are largely unaffected (Fig. 5A, B). These findings are in line with the concept that genes involved in different cellular processes exhibit distinct responses to SOX9 dosage, and SOX9 dosage can underline the severity of a broad spectrum of skeletal abnormalities ^56, 57, 58^.

In addition, we observed an accumulation of IHH and COLX-positive chondrocytes in the *Adgrg6* mutant growth plates, a phenotype not displayed in the *Sox9* cKO mice but was reported in the *Pth1r* cKO mice ^22, 23^. This observation suggests that ADGRG6 may regulate PTHrP/PTH1R/IHH signaling to slow down the progression towards chondrocyte hypertrophy. PTHrP/PTH1R signaling is known to stimulate the Gs/PKA signaling pathway to suppress hypertrophic differentiation by blocking the transcriptional activity of *Runx2* and *Mef2* family members via recruitment of HDAC4^59, 60^. On the other hand, IHH signaling can suppress chondrocyte hypertrophy by stimulating PTHrP expression but also promote hypertrophic differentiation in a PTHrP-independent manner ^7, 61, 62^. *Adgrg6* cKO mice exhibit decreased PTHrP expression in the resting zone but expanded IHH expression in the proliferative zone, suggesting that ADGRG6 helps to regionally refine hypertrophic differentiation by promoting PTHrP expression and by suppressing IHH signaling.

We also revealed a role of ADGRG6 in maintaining chondrocyte survival, as loss of *Adgrg6* leads to increased cell death throughout the growth plates, which mimics the phenotype observed in the *Pth1r* cKO mice and the Gs/Gq double cKO mice but not in the Gs conditional knockout mice ^22, 23^. These data suggest that ADGRG6 may signal through other G protein families, as was recently described for Gi signaling^63^, in addition to Gs. Alternatively, the delayed formation of the secondary ossification center may contribute to the increased cell death in the hypertrophic zone of the *Adgrg6* ckO mice, as the hypertrophic chondrocytes are highly vulnerable to mechanical stress ^64^. In agreement, ablation of *Adgrg6* after the formation of the secondary ossification center (*Adgrg6* cKO^TM^) leads to increased cell death specifically in the resting zone growth plate, supporting the cell-autonomous role of ADGRG6 in maintaining resting chondrocyte homeostasis of the postnatal growth plates.

Previously, we reported that about 50-60% of the *Adgrg6* cKO mice show juvenile-onset scoliosis with a mild IVD histopathology at P20 but exhibit more severe disc degeneration/herniation at 6-8 months ^25, 26, 27^. These observations indicate that the vertebral growth plates may show a delayed response upon loss of *Adgrg6* compared with the growth plates in the long bone. It remains to be determined how ADGRG6/G protein-mediated signaling regulates cell plasticity, survival, and homeostasis of other cartilage populations, such as the articular cartilage.

## Material and Method

### Animal strains

All the animal experiments were approved by the Institutional Animal Care and Use Committee at the University of Southern California (Protocol 21421) and The University of Texas at Austin (Protocol AUP-2021-00215). All mouse strains were described previously, including *Adgrg6^f/f^*(Taconic #TF0269) ^28^, *Col2a1Cre* ^30^, and *AcanCre^ERT2^* ^50^. All mouse strains were kept on a C57BL/6J (JAX:000664) background. Cre (-) littermates were used as wild-type controls. Tamoxifen was administered to control and mutant mice carrying the *AcanCre^ERT2^*allele via intraperitoneal (i.p.) injection from P14 to P18 for 5 consecutive days (1mg/10g body weight, Sigma-Aldrich). Both male and female mice are included in the study.

### Analyses of mice

Skeletal preparations were performed as previously described ^65^. Briefly, P1 pups were skinned and fixed in 10% neutral-buffered formalin for 1 day at 4°C, stained with 0.3% Alcian blue and 0.1% Alizarin red at 37°C for 3 days, and cleared with 1% KOH. Radiographs of the mouse skeleton were generated using a Kubtec DIGIMUS X-ray system (Kubtec T0081B) with auto-exposure under 25 kV as previously described ^25^. The length of the femur is measured on high-resolution radiographic images with ImageJ (https://imagej.nih.gov/ij/download.html).

Histological staining was performed on P1, P14, and P20 knee joints fixed in 10% neutral-buffered formalin (Sigma HT501128) for 1–3 days at room temperature, followed by 1–3 days of decalcification in Formic Acid Bone Decalcifier (Immunocal, StatLab UN3412). After decalcification, tissues were embedded in paraffin and sectioned at 5 mm thickness. Safranin O/Fast Green (SO/FG) staining was performed following standard protocols (Center for Musculoskeletal Research, University of Rochester).

Immunohistochemistry (IHC) or immunofluorescence (IF) analyses were performed on paraffin sections with antigen retrieval (COLII and COLX: 4 mg/ml pepsin in 0.01 N HCl solution, 37°C water bath for 10 min; ADGRG6, SOX9, IHH, GLI2, PRTN3, ITGB3, pCREB, PTH1R, and PTHRP: 10 mM Tris-EDTA with 0.05% Triton-X-100, pH 9.0, 75°C water bath for 5 min; SPP1, MMP9 and MMP13: 100μg/ml hyaluronidase in 1XPBS, 37°C water bath for 10 min; Ki67: 10μg/ml proteinase K in 1XPBS, room temperature for 2 min) and colorimetric developmental methodologies (DAB substrate, Vector Laboratories SK-4105; SG Substrate, Vector Laboratories SK-4700) with the following primary antibodies: anti-GPCR ADGRG6 (Abcam, ab117092, 1:500), anti-SOX9 (EMD Millipore, AB5535, 1:200), anti-type II collagen (COLII) (Thermo Scientific, MS235B, 1:200), anti-type X collagen (COLX) (Abclonal, A18604, 1:100), anti-IHH (Proteintech, 13388-1-AP, 1:100), anti-GLI2 (Santa Cruz Biotech, sc-271786, 1:100), anti-MMP9 (Abclonal, A0289, 1:100), anti-PRTN3 (Abclonal, A19748, 1:100), anti-SPP1 (Cell Signaling, #88742, 1:200), anti-ITGB3 (Abclonal, A2542, 1:100), anti-Phospho-CREB (Ser133) (pCREB) (Cell Signaling, #9198, 1:200), anti-MMP13 (Abcam, ab39012, 1:200), anti-Ki67 (Abcam, ab16667, 1:100), anti-PTH1R (Abclonal, A1744, 1:100) and anti-PTHRP (PTHLH) (Abclonal, A1654, 1:100). Sections were counterstained with hematoxylin or nuclear fast red.

The terminal deoxynucleotidyl transferase dUTP nick-end labeling (TUNEL) cell death assay was performed on paraffin sections with an In Situ Cell Death Detection Kit, Fluorescein (Roche, 11684795910) according to the manufacturer’s instructions. Tissue length and cell number are quantified on the high-resolution bright field or fluorescence images with ImageJ.

### Spatial transcriptomics

Spatial Transcriptomics was conducted using the Visium Spatial Gene Expression System FFPE (formalin-fixed, paraffin-embedded) workflow (10x Genomics). Knee joints of P20 *Col2Cre; Adgrg6^f/f^* (*Adgrg6* cKO) mutants and Cre (-) littermates (wild-type) were harvested and immediately fixed in 10% neutral-buffered formalin (Sigma HT501128) at 4°C for 16 hours with gentle rocking. Samples were thoroughly washed with Milli-Q water (10min x 3 times and 5min x 3 times) at 4°C, and decalcified with Formic Acid Bone Decalcifier (Immunocal, StatLab UN3412) for 16 hours at 4°C with gentle rocking. After decalcification, samples were thoroughly washed with Milli-Q water (10min x 3 times and 5min x 3 times) at 4°C and stored in fresh-made 70% EtOH at 4°C for less than 12 hours. Tissues were processed with a paraffin processor with the following protocol: 45 min x3 with a gradient of EtOH (70%, 80%, 95%); 1h x3 with 100% EtOH; 1h x3 with Xylene; 1h x3 with wax. Samples were immediately embedded after processing. For this project, one control and one mutant knee joint were embedded into a single paraffin block to reduce the batch-to-batch discrepancy. To ensure even chilling and to preserve RNA integrity, the paraffin block was stored in a sealed tissue container at 4°C until needed.

Tissue quality checks and gene expression assays were carried out according to the manufacturer’s protocol. Briefly, slides were deparaffinized, stained with H&E staining protocol, and imaged with a Keyence BZX-710 all-in-one fluorescence microscope (Keyence). After decrosslinking, the Visium mouse transcriptome probe sets were applied to the expression slide, followed by probe release, extension, elution, and library construction. Samples were subjected to pair-ended sequencing using Illumina NovaSeq generating ∼200 M reads.

Alignment was conducted using the Space Ranger pipeline. Subsequent data analysis was conducted using the Loupe Browser (v 6.0.0, 10x Genomics) and Seurat R package (v4.3.0) ^66^. To subset growth plate chondrocytes, spatial spots within the growth plates from the wild-type and *Adgrg6* cKO mice were manually selected using the Loupe Browser. Raw counts of selected growth plate spatial spots were loaded into Seurat using the *Load10X_Spatial* function, and the data were then normalized using the *SCTransform* function (v2, regularization). We integrated the wild-type and Adgrg6 cKO mice datasets using Harmony ^67^ and Seurat Wrappers R packages. To cluster the cells, resolution 0.8 was used for the *FindClusters* function in Seurat. *FindConservedMarkers* function was used to identify conserved chondrocyte populations in the growth plate between wildtype and *Adgrg6* cKO mice, while *FindMarkers* function was used to determine differentially expressed genes within the same chondrocyte population but across two different conditions (wild-type and *Adgrg6* cKO).

Gene Ontology (GO) enrichment analysis of differentially expressed genes was implemented following the instructions provided by the website from The Database for Annotation, Visualization and Integrated Discovery (DAVID) ^68^. Cell-cell crosstalk was analyzed using CellChat R package ^46^.

## Statistics

Statistical analyses were performed in GraphPad Prism 9.5.0 (GraphPad Software Inc, San Diego, CA). Two-tailed Student’s *t*-test was applied as appropriate. A *p*-value of less than 0.05 is considered statistically significant. Bar graphs were generated with mean and standard deviation (SD). At least three biological replicas were analyzed in each experimental group. The number of samples is indicated in the figure legends.

## Data Availability

All the results and analyzed data are available in the main text or the supplementary material. Raw and processed sequencing data will be deposited to GEO.

## Acknowledgments

This work was supported by the National Institute of Arthritis and Musculoskeletal and Skin Diseases (K99/R00AR077090 to Z.L., R00AR075899 and OREF grant to C.L.W., and R01AR072009 to R.S.G.), the ASBMR Fund for Research and Education Research to Z.L., the Start-up Fund to Z.L. from the University of Southern California, and the Faculty Research Seed Grant to Z.L. from the Herman Ostrow School of Dentistry at the University of Southern California. We thank Dr. Fanxin Long for sharing the *Col2a1Cre* mouse strain.

## Author Contributions

Conceptualization: Z.L., R.S.G, and C.L.W. Methodology: F.B., V.H., H.C.F., Y.C., R.S.G, C.L.W. and Z.L. Investigation: F.B., V.H., H.C.F., Y.C., C.L.W, and Z.L. Visualization: F.B., V.H., H.C.F., Y.C., C.L.W, and Z.L. Funding acquisition: Z.L., R.S.G, and C.L.W. Project administration: Z.L. Supervision: Z.L. and C.L.W. Writing-original draft: F.B. and Z.L. Writing-review & editing: F.B., V.H., H.C.F., Y.C., R.S.G, C.L.W. and Z.L.

---

**Supplementary Figure 1.**
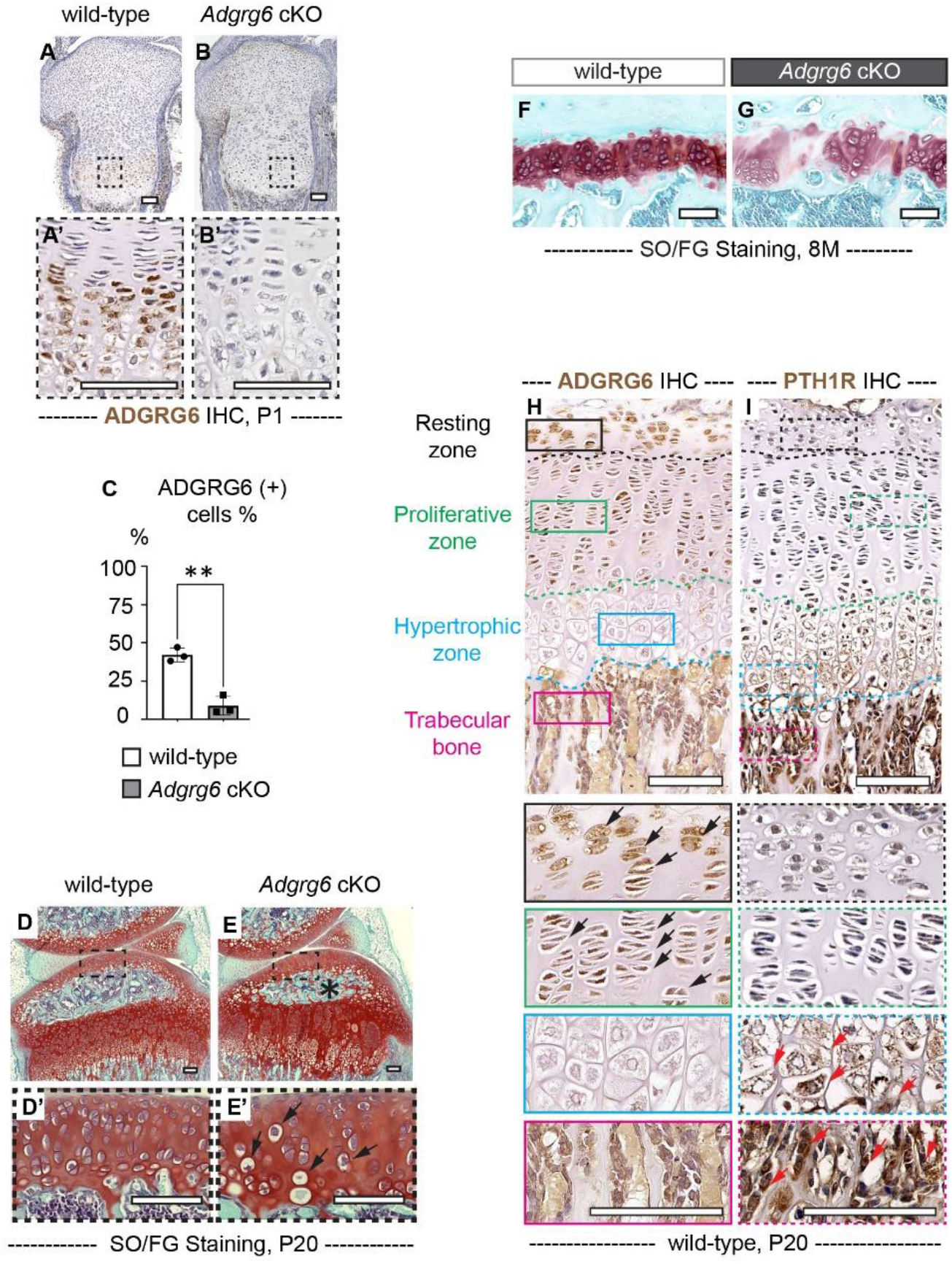
ADGRG6 is required in the homeostasis of the growth plate. **(A-C)** IHC analyses reveal reduced ADGRG6 expression in *Adgrg6* cKO mutants (B, B’) compared with the controls (A, A’) at P1. The percentage of ADGRG6 (+) cells is quantified in C. (n=3 for each genotype. **p= 0.0017.) Bars are plotted by means ± SD, two-tailed *t*-test. **(D-E’)** Loss of *Adgrg6* results in reduced cellularity and abnormal hypertrophic differentiation in articular cartilage (D’ and E’). (n=7 per genotype.) Black arrows in E’ indicate hypertrophic-like chondrocytes. Panel D and E are the same panel as shown in Figure. 1H, I. **(F, G)** The SO/FG staining of 8-month-old control and mutant growth plates. (n=4 per genotype.) **(H, I)** IHC analyses of ADGRG6 and PTH1R of P20 wild-type mice. Higher magnification of the resting zone, proliferative one, hypertrophic zone, and trabecular bone are shown in black, green, blue, and magenta boxes. ADGRG6 (+) and PTH1R (+) cells are indicated with black and red arrows, respectively. (n=3 for each genotype.) Panel F is the same panel as shown in Figure 1J. SO/FG: Safranin O/Fast Green. IHC: Immunohistochemistry. Scale bar: 100µm.

**Supplementary Figure 2.**
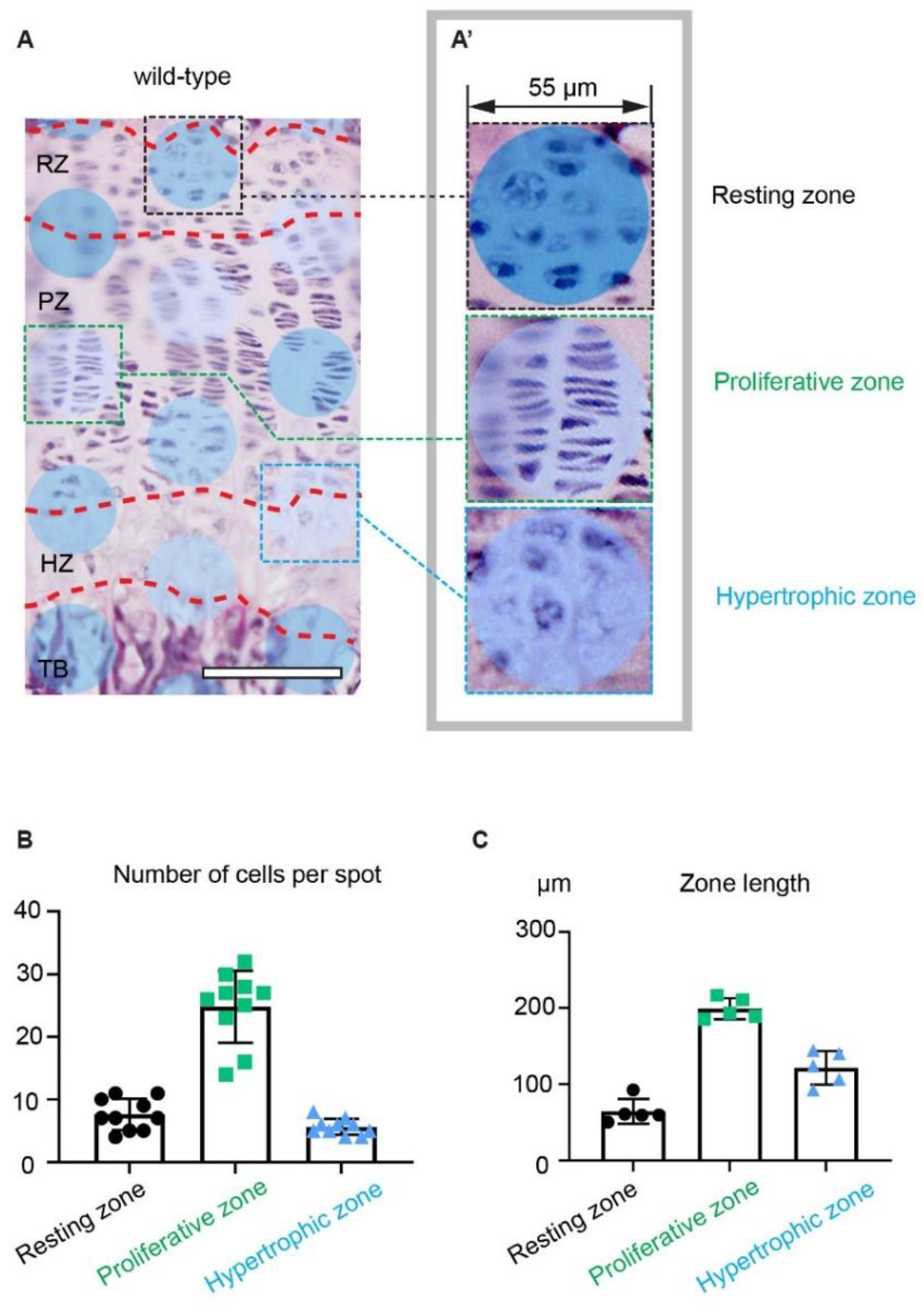
One Visium capture spot can cover multiple cells in the growth plate. **(A, A’)** Overlay of the Visium capture spot on top of the H&E-stained sections. Representative spots mapped to the resting zone, proliferative zone, and hypertrophic zone are shown in A’. Contrast for images in A’ was enhanced with Adobe Photoshop to better show the nuclei. Images were exported from the Loupe Browser. **(B)** Quantification of the number of cells covered by each capture spot (n=10 spots per zone.) **(C)** Quantification of the zone length of the wild-type growth plate. (n=5 mice.) Scale bar: 100µm.

**Supplementary Figure 3.**
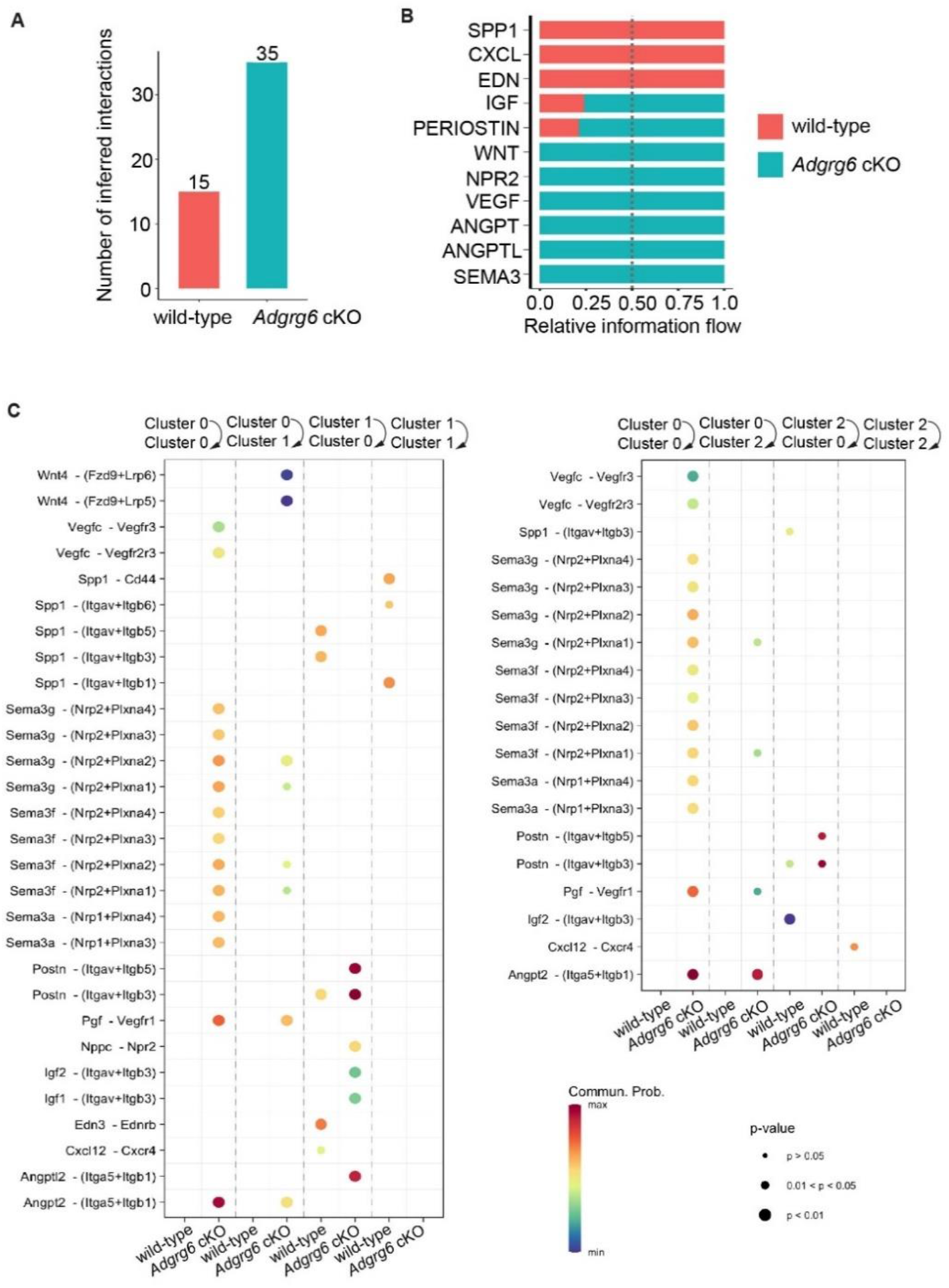
CellChat analyses revealed altered cell-cell crosstalk in the mutant growth plate. **(A, B)** The number of inferred interactions (A) and relative information flow (B) are shown in the control and the mutant mice. **(C)** Dot plots show all the significant ligand-receptor pairs that involve in the crosstalk between and within cluster 0 and cluster 1, and cluster 0 and cluster 2. The dot color and size represent the calculated communication probability and p-values. p-values are computed from the one-side permutation test.

**Supplementary Figure 4.**
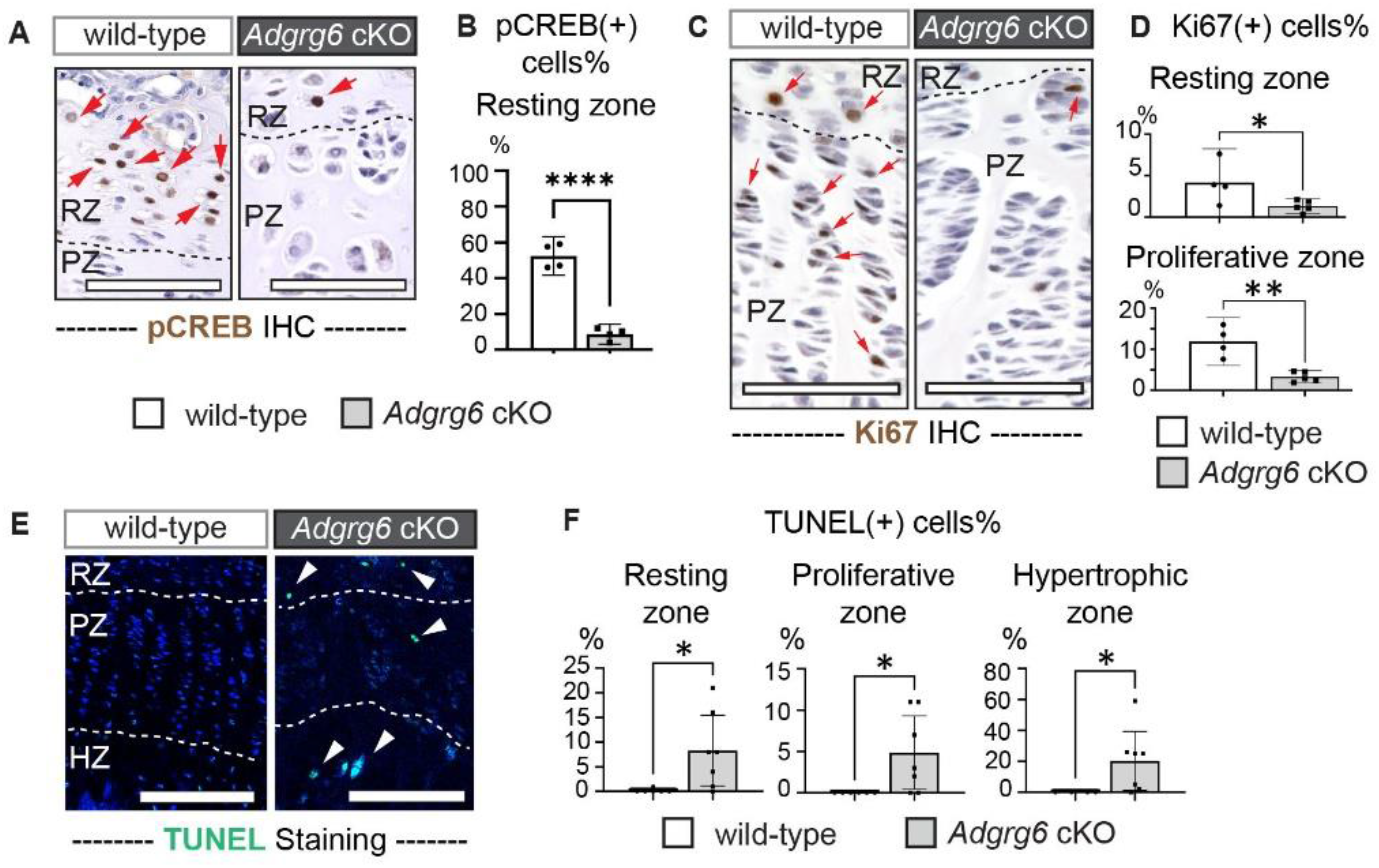
ADGRG6 maintains growth plate cell proliferation and survival. (A, B) IHC analysis reveals reduced pCREB expression in the resting zone growth plate of the *Adgrg6* cKO mutants at P20. (n=4 per genotype. ****p<0.0001.) (C, D) IHC analysis reveals reduced Ki67 expression in the resting proliferative zones of the growth plate in *Adgrg6* cKO mutants at P20. (n=4 for wild-type, n=5 for *Adgrg6* cKO. *p= 0.0482, **p= 0.0016.) (E, F) TUNEL staining reveals increased cell death in all the zones of the mutant growth plate at P20. (n=6 for wild-type, n=7 for *Adgrg6* cKO. F: *p= 0.0273 for resting zone; *p= 0.0318 for proliferative zone; *p= 0.0383 for hypertrophic zone.) Bars are plotted by means ± SD, two-tailed *t*-test. Scale bar: 100µm.

**Supplementary Figure 5.**
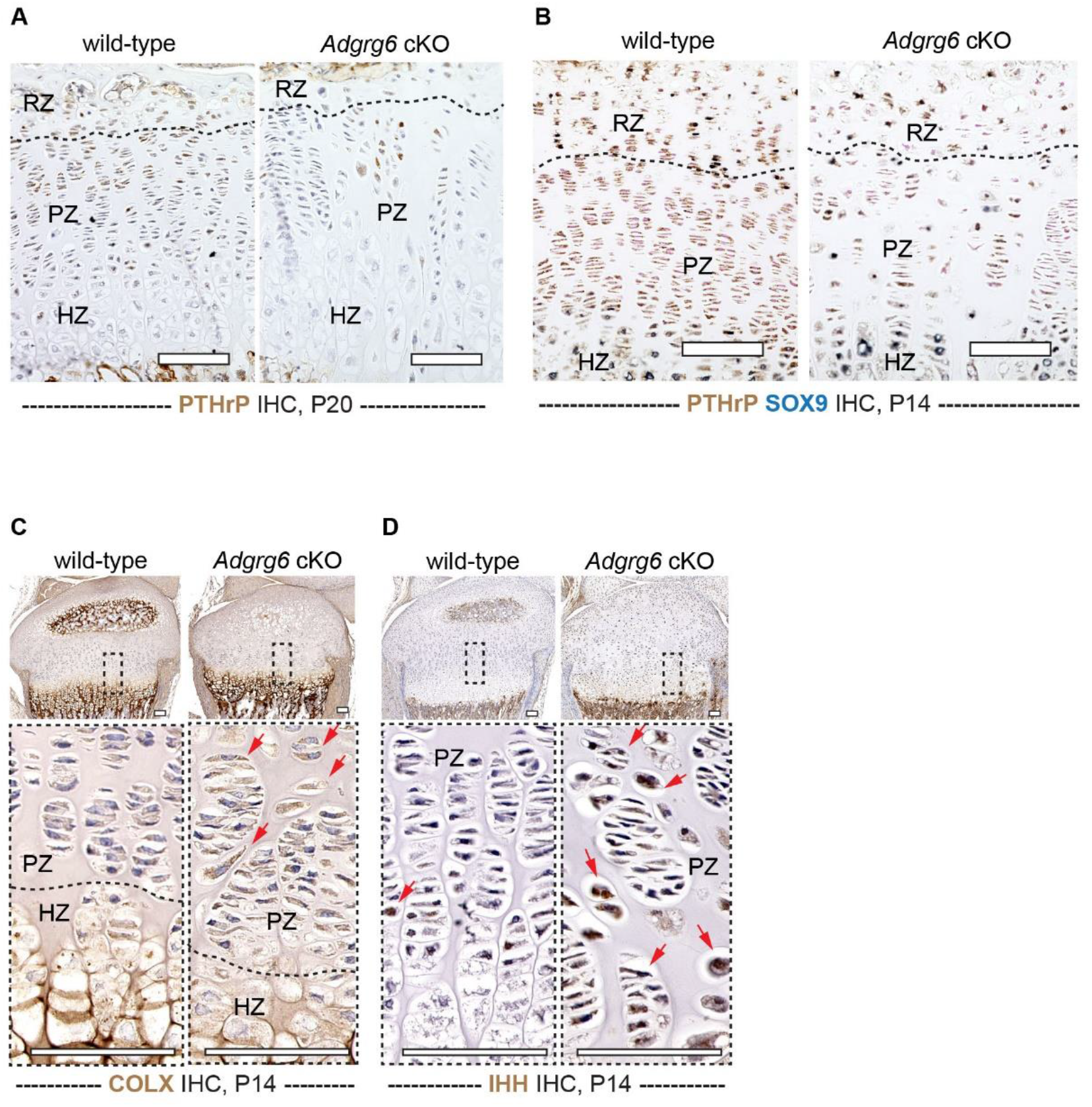
ADGRG6 is required for the proper formation of the growth plate. (A) IHC analysis revealed reduced expression of PTHrP in P20 *Adgrg6* cKO mutant growth plates. (n=3 per genotype.) (B) Large-scale images of dual color IHC analysis of PTHrP and SOX9 as shown in Figure 6 M, N. (n=3 per genotype.) (C, D) IHC analyses reveal diffused expression of COLX (C) and IHH (D) in mutant growth plates at P14. COLX (+) and IHH (+) cells are indicated with red arrows in C and D. (n=3 per genotype.) IHC: Immunohistochemistry. RZ: resting zone; PZ: proliferative zone; HZ: hypertrophic zone. Scale bar: 100µm.

**Supplementary Figure 6.**
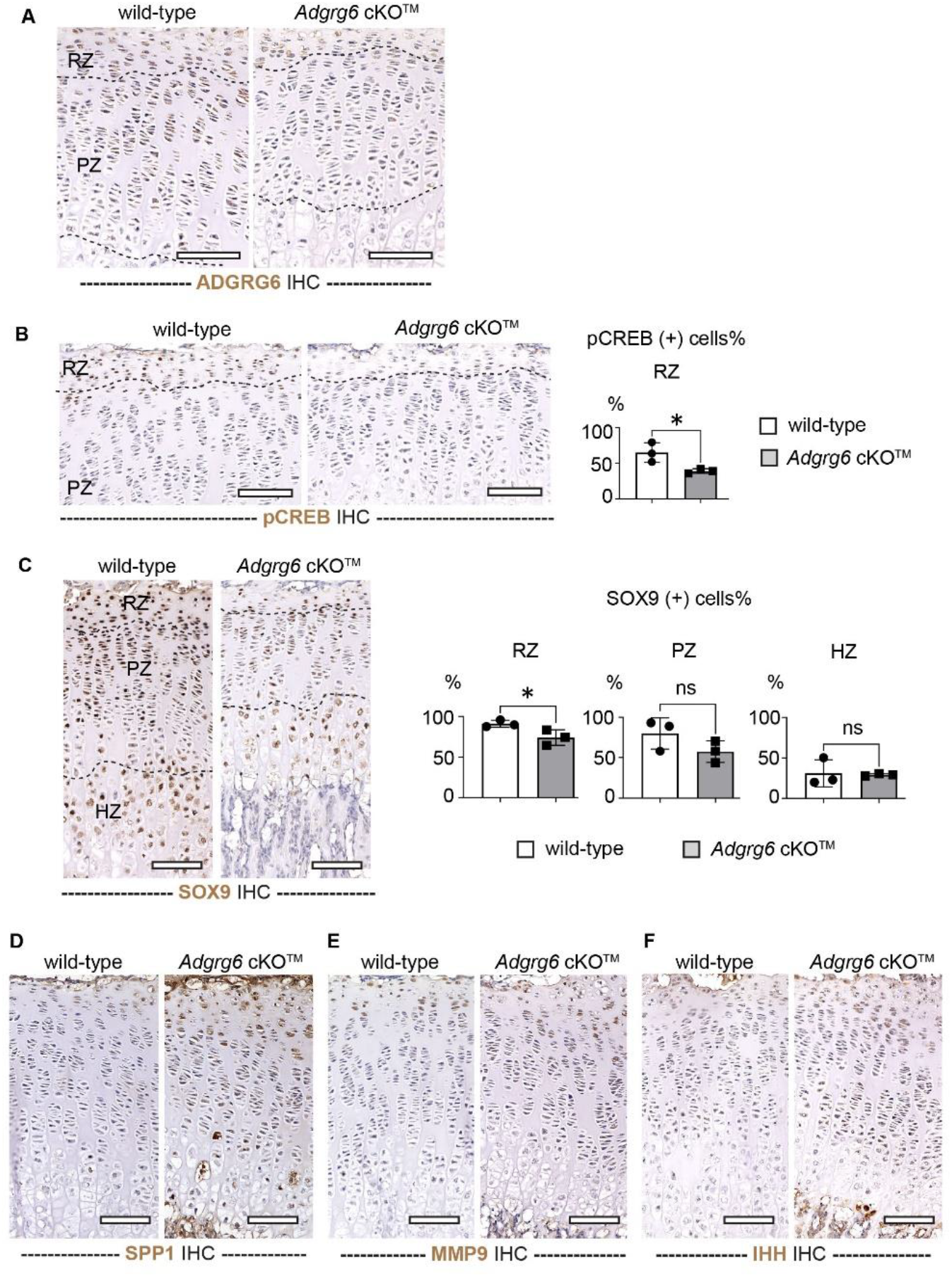
ADGRG6 is required for postnatal growth plate homeostasis. **(A, B)** IHC analysis shows a reduced expression of ADGRG6 and pCREB in the growth plate of the *Adgrg6* cKO^TM^ mutant mice at P20. Panel A and B are the large-scale images as shown in Figure 7J, K. (n=3 per genotype. *p= 0.0329.) **(C)** IHC analysis shows a decreased expression of SOX9 in the resting zone of the *Adgrg6* cKO^TM^ mutant mice at P20. Large-scale images as shown in Figure 7L. (n=3 per genotype. *p= 0.0472 in RZ; ns, p= 0.8458 in PZ; ns, p= 0.1700 in HZ.) Bars are plotted by means ± SD, two-tailed *t*-test. **(D-F)** Large-scale images of IHC analyses of SPP1 (D), MMP9 (E), and IHH (F) as shown in Figure 7 M, N, and O. IHC: Immunohistochemistry. RZ: resting zone; PZ: proliferative zone; HZ: hypertrophic zone. Scale bar: 100µm.

## References

1. Kronenberg HM. Developmental regulation of the growth plate. Nature 423, 332–336 (2003).

2. Melrose J, Shu C, Whitelock JM, Lord MS. The cartilage extracellular matrix as a transient developmental scaffold for growth plate maturation. Matrix Biol 52-54, 363-383 (2016).

3. Krishnan Y, Grodzinsky AJ. Cartilage diseases. Matrix Biol 71-72, 51–69 (2018).

4. Mizuhashi K, et al. Resting zone of the growth plate houses a unique class of skeletal stem cells. Nature 563, 254–258 (2018).

5. Newton PT, et al. A radical switch in clonality reveals a stem cell niche in the epiphyseal growth plate. Nature 567, 234–238 (2019).

6. Schipani E, Provot S. PTHrP, PTH, and the PTH/PTHrP receptor in endochondral bone development. Birth Defects Res C Embryo Today 69, 352–362 (2003).

7. Ohba S. Hedgehog Signaling in Endochondral Ossification. J Dev Biol 4, (2016).

8. Yang L, Tsang KY, Tang HC, Chan D, Cheah KS. Hypertrophic chondrocytes can become osteoblasts and osteocytes in endochondral bone formation. Proc Natl Acad Sci U S A 111, 12097–12102 (2014).

9. Zhou X, von der Mark K, Henry S, Norton W, Adams H, de Crombrugghe B. Chondrocytes transdifferentiate into osteoblasts in endochondral bone during development, postnatal growth and fracture healing in mice. PLoS Genet 10, e1004820 (2014).

10. Long JT, et al. Hypertrophic chondrocytes serve as a reservoir for marrow-associated skeletal stem and progenitor cells, osteoblasts, and adipocytes during skeletal development. Elife 11, (2022).

11. Haseeb A, et al. SOX9 keeps growth plates and articular cartilage healthy by inhibiting chondrocyte dedifferentiation/osteoblastic redifferentiation. Proc Natl Acad Sci U S A 118, (2021).

12. Sriram K, Insel PA. G Protein-Coupled Receptors as Targets for Approved Drugs: How Many Targets and How Many Drugs? Mol Pharmacol 93, 251–258 (2018).

13. Weis WI, Kobilka BK. The Molecular Basis of G Protein-Coupled Receptor Activation. Annu Rev Biochem 87, 897–919 (2018).

14. Wootten D, Christopoulos A, Marti-Solano M, Babu MM, Sexton PM. Mechanisms of signalling and biased agonism in G protein-coupled receptors. Nat Rev Mol Cell Biol 19, 638–653 (2018).

15. Syrovatkina V, Alegre KO, Dey R, Huang XY. Regulation, Signaling, and Physiological Functions of G-Proteins. J Mol Biol 428, 3850–3868 (2016).

16. Amizuka N, Warshawsky H, Henderson JE, Goltzman D, Karaplis AC. Parathyroid hormone-related peptide-depleted mice show abnormal epiphyseal cartilage development and altered endochondral bone formation. J Cell Biol 126, 1611–1623 (1994).

17. Karaplis AC, et al. Lethal skeletal dysplasia from targeted disruption of the parathyroid hormone-related peptide gene. Genes Dev 8, 277–289 (1994).

18. Lanske B, et al. PTH/PTHrP receptor in early development and Indian hedgehog-regulated bone growth. Science 273, 663–666 (1996).

19. Miao D, He B, Karaplis AC, Goltzman D. Parathyroid hormone is essential for normal fetal bone formation. J Clin Invest 109, 1173–1182 (2002).

20. Kobayashi T, et al. PTHrP and Indian hedgehog control differentiation of growth plate chondrocytes at multiple steps. Development 129, 2977–2986 (2002).

21. Amano K, Densmore M, Fan Y, Lanske B. Ihh and PTH1R signaling in limb mesenchyme is required for proper segmentation and subsequent formation and growth of digit bones. Bone 83, 256–266 (2016).

22. Hirai T, Chagin AS, Kobayashi T, Mackem S, Kronenberg HM. Parathyroid hormone/parathyroid hormone-related protein receptor signaling is required for maintenance of the growth plate in postnatal life. Proc Natl Acad Sci U S A 108, 191–196 (2011).

23. Chagin AS, et al. G-protein stimulatory subunit alpha and Gq/11alpha G-proteins are both required to maintain quiescent stem-like chondrocytes. Nat Commun 5, 3673 (2014).

24. Chagin AS, Kronenberg HM. Role of G-proteins in the differentiation of epiphyseal chondrocytes. J Mol Endocrinol 53, R39–45 (2014).

25. Liu Z, et al. An adhesion G protein-coupled receptor is required in cartilaginous and dense connective tissues to maintain spine alignment. Elife 10, (2021).

26. Liu Z, et al. Dysregulation of STAT3 signaling is associated with endplate-oriented herniations of the intervertebral disc in Adgrg6 mutant mice. PLoS Genet 15, e1008096 (2019).

27. Karner CM, Long F, Solnica-Krezel L, Monk KR, Gray RS. Gpr126/Adgrg6 deletion in cartilage models idiopathic scoliosis and pectus excavatum in mice. Hum Mol Genet 24, 4365–4373 (2015).

28. Mogha A, et al. Gpr126 functions in Schwann cells to control differentiation and myelination via G-protein activation. J Neurosci 33, 17976–17985 (2013).

29. Hai P. Nguyen AU, Rory Sheng, Cassidy Biellak, Kelly An, Hélène Choquet,, Thomas J. Hoffman RSG, Nadav Ahituv. ADGRG6 promotes adipogenesis and is involved in sex-specific fat distribution. bioRxiv preprint (2022).

30. Long F, Zhang XM, Karp S, Yang Y, McMahon AP. Genetic manipulation of hedgehog signaling in the endochondral skeleton reveals a direct role in the regulation of chondrocyte proliferation. Development 128, 5099–5108 (2001).

31. Si J, Wang C, Zhang D, Wang B, Zhou Y. Osteopontin in Bone Metabolism and Bone Diseases. Med Sci Monit 26, e919159 (2020).

32. Lefebvre V, Angelozzi M, Haseeb A. SOX9 in cartilage development and disease. Curr Opin Cell Biol 61, 39–47 (2019).

33. Miles RR, et al. ADAMTS-1: A cellular disintegrin and metalloprotease with thrombospondin motifs is a target for parathyroid hormone in bone. Endocrinology 141, 4533–4542 (2000).

34. Sabik OL, Calabrese GM, Taleghani E, Ackert-Bicknell CL, Farber CR. Identification of a Core Module for Bone Mineral Density through the Integration of a Co-expression Network and GWAS Data. Cell Rep 32, 108145 (2020).

35. Komori T. Molecular Mechanism of Runx2-Dependent Bone Development. Mol Cells 43, 168–175 (2020).

36. Chan CK, et al. Clonal precursor of bone, cartilage, and hematopoietic niche stromal cells. Proc Natl Acad Sci U S A 110, 12643–12648 (2013).

37. Deng C, Wynshaw-Boris A, Zhou F, Kuo A, Leder P. Fibroblast growth factor receptor 3 is a negative regulator of bone growth. Cell 84, 911–921 (1996).

38. Swahn H, et al. Senescent cell population with ZEB1 transcription factor as its main regulator promotes osteoarthritis in cartilage and meniscus. Ann Rheum Dis 82, 403–415 (2023).

39. Goldring MB, Tsuchimochi K, Ijiri K. The control of chondrogenesis. J Cell Biochem 97, 33–44 (2006).

40. Moses MA, et al. Troponin I is present in human cartilage and inhibits angiogenesis. Proc Natl Acad Sci U S A 96, 2645–2650 (1999).

41. Zhu X, et al. A gain-of-function mutation in Tnni2 impeded bone development through increasing Hif3a expression in DA2B mice. PLoS Genet 10, e1004589 (2014).

42. Palmer GD, et al. F-spondin deficient mice have a high bone mass phenotype. PLoS One 9, e98388 (2014).

43. Pekkinen M, et al. Osteoporosis and skeletal dysplasia caused by pathogenic variants in SGMS2. JCI Insight 4, (2019).

44. Deng Z, et al. Temporal transcriptome features identify early skeletal commitment during human epiphysis development at single-cell resolution. iScience 26, 107200 (2023).

45. Eyre DR, Weis MA, Wu JJ. Articular cartilage collagen: an irreplaceable framework? Eur Cell Mater 12, 57–63 (2006).

46. Jin S, et al. Inference and analysis of cell-cell communication using CellChat. Nat Commun 12, 1088 (2021).

47. Bonnet N, Conway SJ, Ferrari SL. Regulation of beta catenin signaling and parathyroid hormone anabolic effects in bone by the matricellular protein periostin. Proc Natl Acad Sci U S A 109, 15048–15053 (2012).

48. Alto LT, Terman JR. Semaphorins and their Signaling Mechanisms. Methods Mol Biol 1493, 1–25 (2017).

49. Guasto A, Cormier-Daire V. Signaling Pathways in Bone Development and Their Related Skeletal Dysplasia. Int J Mol Sci 22, (2021).

50. Henry SP, Jang CW, Deng JM, Zhang Z, Behringer RR, de Crombrugghe B. Generation of aggrecan-CreERT2 knockin mice for inducible Cre activity in adult cartilage. Genesis 47, 805–814 (2009).

51. Henry SP, Liang S, Akdemir KC, de Crombrugghe B. The postnatal role of Sox9 in cartilage. J Bone Miner Res 27, 2511–2525 (2012).

52. Piera-Velazquez S, Hawkins DF, Whitecavage MK, Colter DC, Stokes DG, Jimenez SA. Regulation of the human SOX9 promoter by Sp1 and CREB. Exp Cell Res 313, 1069–1079 (2007).

53. Huang W, Zhou X, Lefebvre V, de Crombrugghe B. Phosphorylation of SOX9 by cyclic AMP-dependent protein kinase A enhances SOX9’s ability to transactivate a Col2a1 chondrocyte-specific enhancer. Mol Cell Biol 20, 4149–4158 (2000).

54. Huang W, Chung UI, Kronenberg HM, de Crombrugghe B. The chondrogenic transcription factor Sox9 is a target of signaling by the parathyroid hormone-related peptide in the growth plate of endochondral bones. Proc Natl Acad Sci U S A 98, 160–165 (2001).

55. Zhao L, Li G, Zhou GQ. SOX9 directly binds CREB as a novel synergism with the PKA pathway in BMP-2-induced osteochondrogenic differentiation. J Bone Miner Res 24, 826–836 (2009).

56. Naqvi S, et al. Precise modulation of transcription factor levels identifies features underlying dosage sensitivity. Nat Genet 55, 841–851 (2023).

57. Au TYK, et al. Hypomorphic and dominant-negative impact of truncated SOX9 dysregulates Hedgehog-Wnt signaling, causing campomelia. Proc Natl Acad Sci U S A 120, e2208623119 (2023).

58. Wang L, et al. Variants in the SOX9 transactivation middle domain induce axial skeleton dysplasia and scoliosis. medRxiv preprint, (2023).

59. Kozhemyakina E, Cohen T, Yao TP, Lassar AB. Parathyroid hormone-related peptide represses chondrocyte hypertrophy through a protein phosphatase 2A/histone deacetylase 4/MEF2 pathway. Mol Cell Biol 29, 5751–5762 (2009).

60. Correa D, et al. Zfp521 is a target gene and key effector of parathyroid hormone-related peptide signaling in growth plate chondrocytes. Dev Cell 19, 533–546 (2010).

61. Karp SJ, Schipani E, St-Jacques B, Hunzelman J, Kronenberg H, McMahon AP. Indian hedgehog coordinates endochondral bone growth and morphogenesis via parathyroid hormone related-protein-dependent and -independent pathways. Development 127, 543–548 (2000).

62. Mak KK, Kronenberg HM, Chuang PT, Mackem S, Yang Y. Indian hedgehog signals independently of PTHrP to promote chondrocyte hypertrophy. Development 135, 1947–1956 (2008).

63. Wilde C, Chaudhry PM, Luo R, Simon KU, Piao X, Liebscher I. Collagen VI Is a Gi-Biased Ligand of the Adhesion GPCR GPR126/ADGRG6. Cells 12, (2023).

64. Xie M, et al. Secondary ossification center induces and protects growth plate structure. Elife 9, (2020).

65. Liu Z, Ramachandran J, Vokes SA, Gray RS. Regulation of terminal hypertrophic chondrocyte differentiation in Prmt5 mutant mice modeling infantile idiopathic scoliosis. Dis Model Mech 12, (2019).

66. Hao Y, et al. Integrated analysis of multimodal single-cell data. Cell 184, 3573–3587 e3529 (2021).

67. Korsunsky I, et al. Fast, sensitive and accurate integration of single-cell data with Harmony. Nat Methods 16, 1289–1296 (2019).

68. Sherman BT, et al. DAVID: a web server for functional enrichment analysis and functional annotation of gene lists (2021 update). Nucleic Acids Res 50, W216–W221 (2022).

