## Supplementary figure for "Spatial Transcriptomics Reveals the Requirement of ADGRG6 in Maintaining Chondrocyte Homeostasis in Mouse Growth Plates"

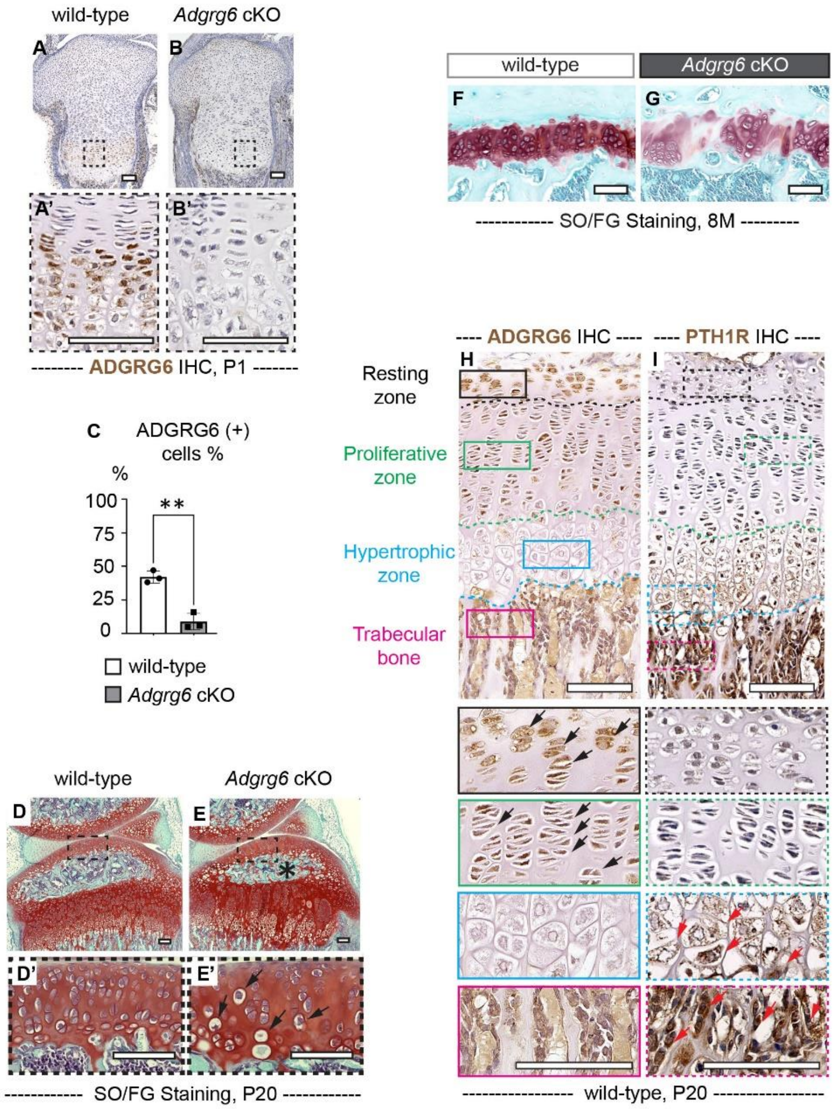


**Supplementary Figure 1. ADGRG6 is required in the homeostasis of the growth plate. (A-C)** IHC analyses reveal reduced ADGRG6 expression in *Adgrg6* cKO mutants (B, B') compared with the controls (A, A') at P1. The percentage of ADGRG6 (+) cells is quantified in C. (n=3 for each genotype. **p= 0.0017.) Bars are plotted by means ± SD, two-tailed *t*-test. **(D-E')** Loss of *Adgrg6* results in reduced cellularity and abnormal hypertrophic differentiation in articular cartilage (D' and E'). (n=7 per genotype.) Black arrows in E' indicate hypertrophic-like chondrocytes. Panel D and E are the same panel as shown in Figure. 1H, I. **(F, G)** The SO/FG staining of 8-month-old control and mutant growth plates. (n=4 per genotype.) **(H, I)** IHC analyses of ADGRG6 and PTH1R of P20 wild-type mice. Higher magnification of the resting zone, proliferative one, hypertrophic zone, and trabecular bone are shown in black, green, blue, and magenta boxes. ADGRG6 (+) and PTH1R (+) cells are indicated with black and red arrows, respectively. (n=3 for each genotype.) Panel F is the same panel as shown in Figure 1J. SO/FG: Safranin O/Fast Green. IHC: Immunohistochemistry. Scale bar: 100μm.

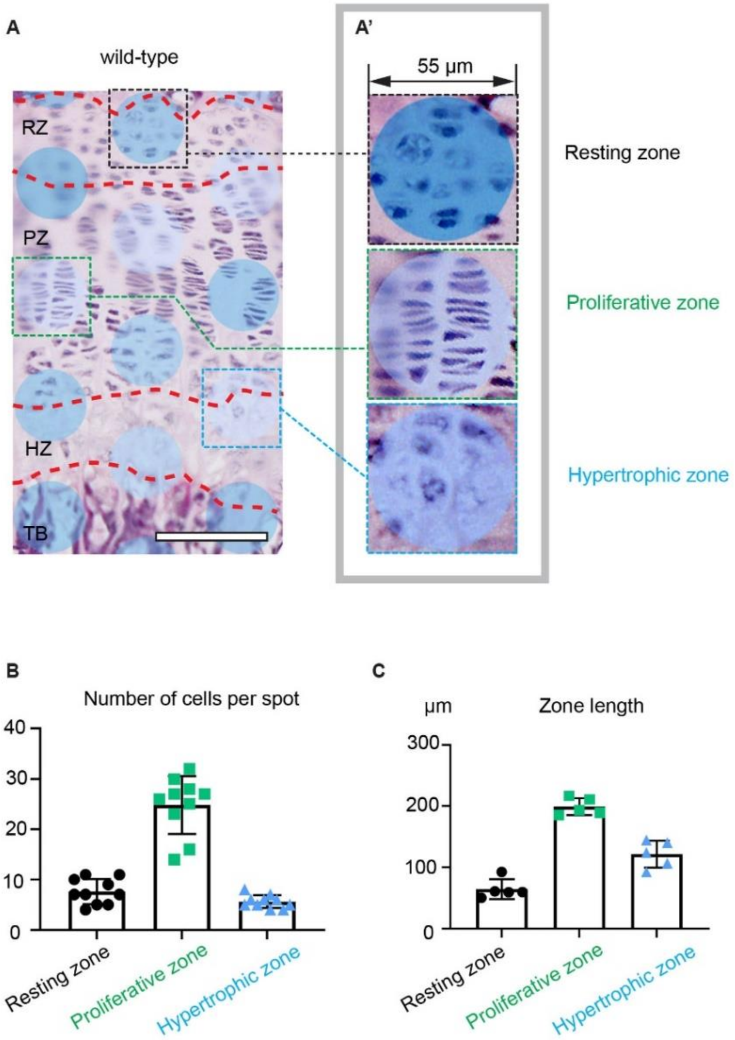


**Supplementary Figure 2. One Visium capture spot can cover multiple cells in the growth plate. (A, A')** Overlay of the Visium capture spot on top of the H&E-stained sections. Representative spots mapped to the resting zone, proliferative zone, and hypertrophic zone are shown in A'. Contrast for images in A' was enhanced with Adobe Photoshop to better show the nuclei. Images were exported from the Loupe Browser. **(B)** Quantification of the number of cells covered by each capture spot (n=10 spots per zone.) **(C)** Quantification of the zone length of the wild-type growth plate. (n=5 mice.) Scale bar: 100µm.

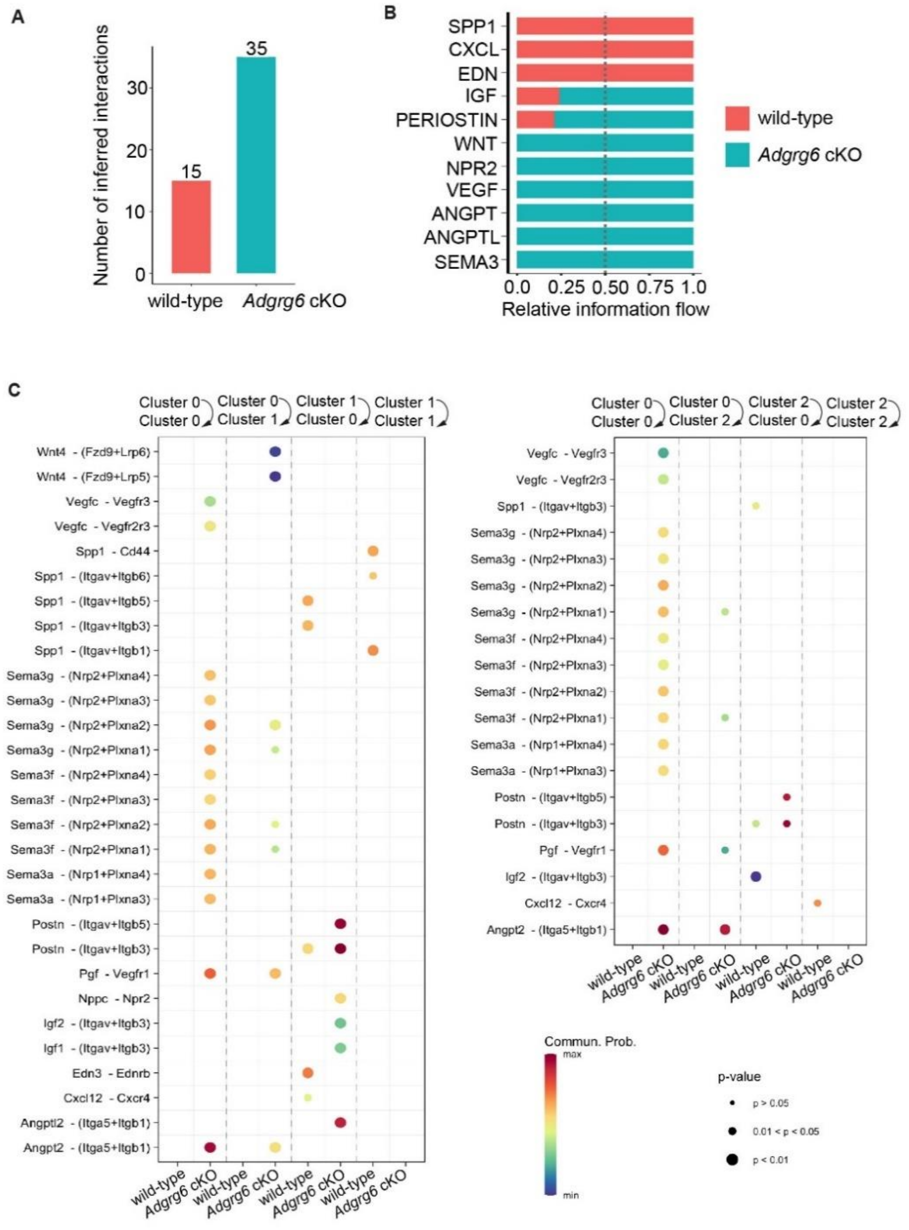


**Supplementary Figure 3. CellChat analyses revealed altered cell-cell crosstalk in the mutant growth plate. (A, B)** The number of inferred interactions (A) and relative information flow (B) are shown in the control and the mutant mice. **(C)** Dot plots show all the significant ligand-receptor pairs that involve in the crosstalk between and within cluster 0 and cluster 1, and cluster 0 and cluster 2. The dot color and size represent the calculated communication probability and p-values. p-values are computed from the one-side permutation test.

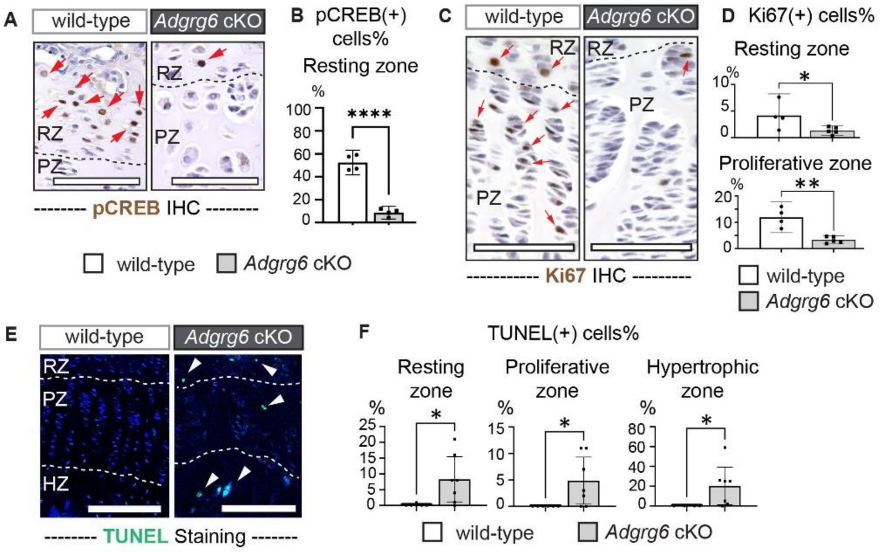


**Supplementary Figure 4. ADGRG6 maintains growth plate cell proliferation and survival.** **(A, B)** IHC analysis reveals reduced pCREB expression in the resting zone growth plate of the *Adgrg6* cKO mutants at P20. (n=4 per genotype. ****p<0.0001.) (**C, D**) IHC analysis reveals reduced Ki67 expression in the resting proliferative zones of the growth plate in *Adgrg6* cKO mutants at P20. (n=4 for wild -type, n=5 for *Adgrg6* cKO. *p= 0.0482, **p= 0.0016.) **(E, F)** TUNEL staining reveals increased cell death in all the zones of the mutant growth plate at P20. (n=6 for wild -type, n=7 for *Adgrg6* cKO. F: *p= 0.0273 for resting zone; *p= 0.0318 for proliferative zone; *p= 0.0383 for hypertrophic zone.) Bars are plotted by means ± SD, two-tailed *t*-test. Scale bar: 100μm.

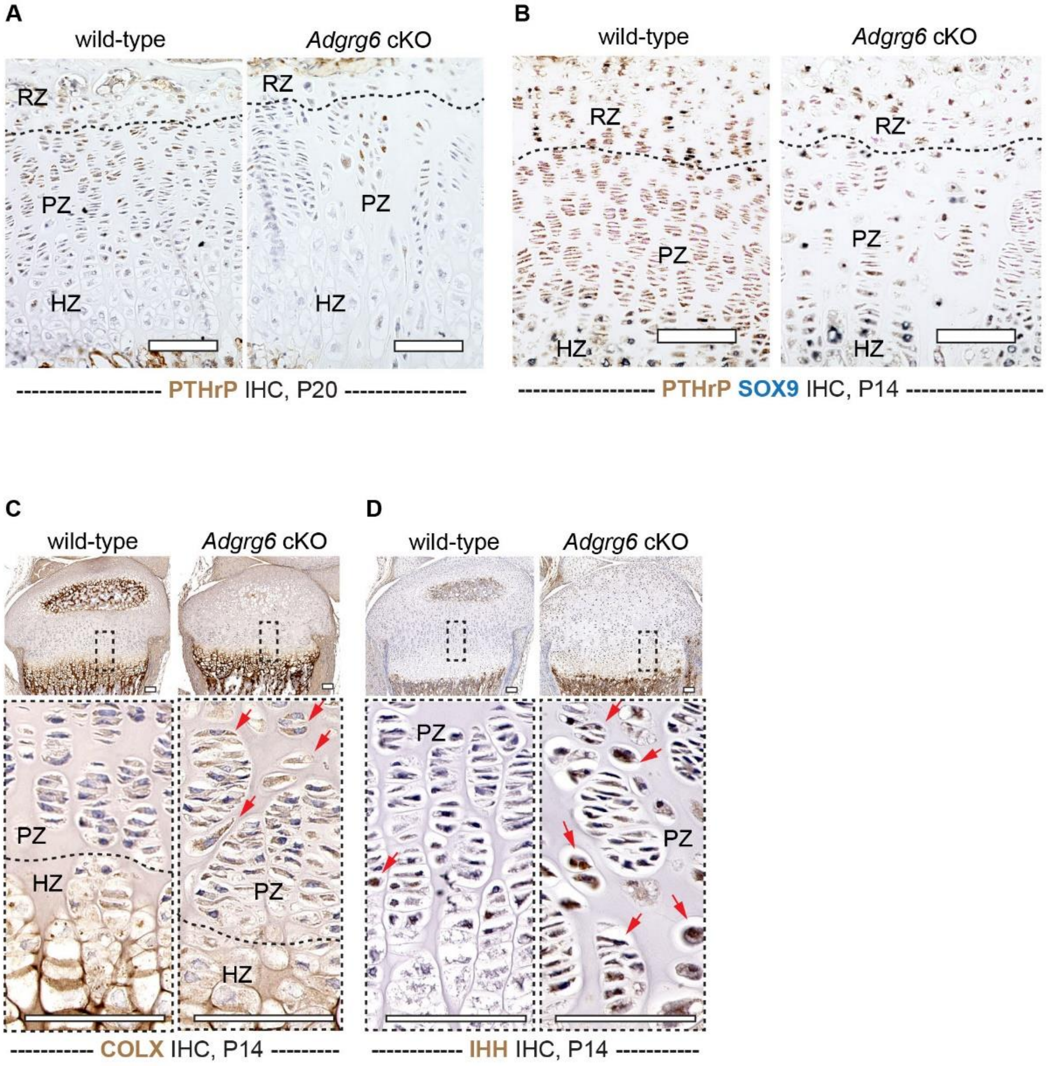


**Supplementary Figure 5. ADGRG6 is required for the proper formation of the growth plate. (A)** IHC analysis revealed reduced expression of PTHrP in P20 *Adgrg6* cKO mutant growth plates. (n=3 per genotype.) **(B)** Large-scale images of dual color IHC analysis of PTHrP and SOX9 as shown in Figure 6 M, N. (n=3 per genotype.) (**C, D**) IHC analyses reveal diffused expression of COLX (C) and IHH (D) in mutant growth plates at P14. COLX (+) and IHH (+) cells are indicated with red arrows in C and D. (n=3 per genotype.) IHC: Immunohistochemistry. RZ: resting zone; PZ: proliferative zone; HZ: hypertrophic zone. Scale bar: 100μm.

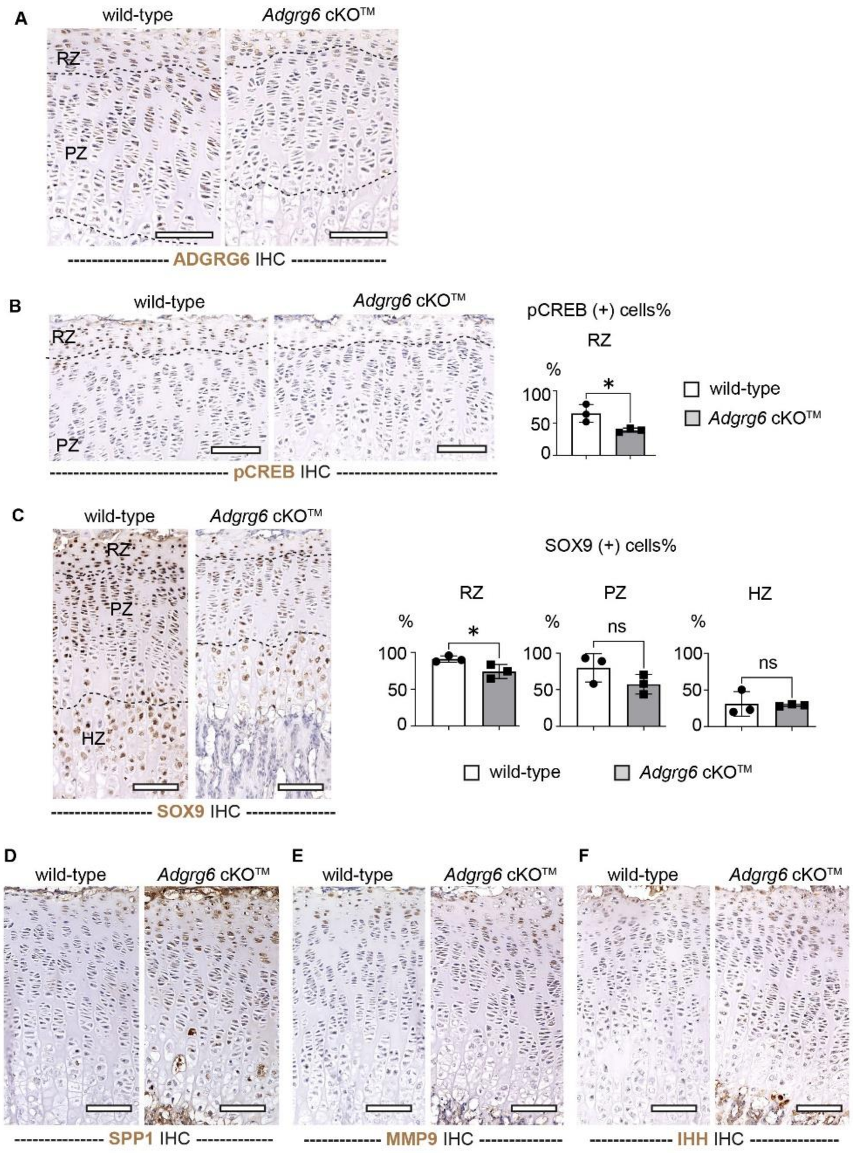


**Supplementary Figure 6. ADGRG6 is required for postnatal growth plate homeostasis. (A, B)** IHC analysis shows a reduced expression of ADGRG6 and pCREB in the growth plate of the *Adgrg6* cKO$^{TM}$ mutant mice at P20. Panel A and B are the large-scale images as shown in Figure 7J, K. (n=3 per genotype. *p= 0.0329.) **(C)** IHC analysis shows a decreased expression of SOX9 in the resting zone of the *Adgrg6* cKO$^{TM}$ mutant mice at P20. Large-scale images as shown in Figure 7L. (n=3 per genotype. *p= 0.0472 in RZ; ns, p= 0.8458 in PZ; ns, p= 0.1700 in HZ.) Bars are plotted by means ± SD, two-tailed *t*-test. **(D-F)** Large-scale images of IHC analyses of SPP1 (D), MMP9 (E), and IHH (F) as shown in Figure 7 M, N, and O. IHC: Immunohistochemistry. RZ: resting zone; PZ: proliferative zone; HZ: hypertrophic zone. Scale bar: 100μm.
